# Structural characterization of stem cell factors Oct4, Sox2, Nanog and Esrrb disordered domains, and a method to identify their phospho-dependent binding partners

**DOI:** 10.1101/2023.03.05.531149

**Authors:** Bouguechtouli Chafiaa, Rania Ghouil, Ania Alik, Dingli Florent, Loew Damarys, Theillet Francois-Xavier

## Abstract

The combined expression of a handful of pluripotency transcription factors (PluriTFs) in somatic cells can generate induced pluripotent stem cells (iPSCs). Here, we report the structural characterization of disordered regions contained in four important PluriTFs, namely Oct4, Sox2, Nanog and Esrrb. Moreover, many post-translational modifications (PTMs) have been detected on PluriTFs, whose roles are not yet characterized. To help in their study, we also present a method i) to produce well-characterized phosphorylation states of PluriTFs, using NMR analysis, and ii) to use them for pull-downs in stem cell extracts analyzed by quantitative proteomics to identify of Sox2 binders.

## 1. Introduction

The possibility of reprogramming somatic cells to an induced pluripotency state was revealed in the 2000s, carrying great expectations in the fields of Biology and Medicine [1–3]. Induced pluripotent stem cells (iPSCs) and embryonic stem cells (ESCs) are characterized by the active state of a pluripotency network, whose core comprises the pluripotency transcription factors (PluriTFs) Oct4, Sox2, Nanog and Esrrb (OSNE). These bind to enhancer sequences, and thus activate or repress, or even “bookmark” during mitosis, a wealth of genes related to pluripotency or cell differentiation [4–9]. Consistently, their misregulation correlates with cancer malignancy and stemness [10–14].

Comprehensive structural descriptions of OSNE are missing to our knowledge. The folded DNA-binding domains (DBDs) of OSNE have been structurally characterized, in complex with their DNA target sequences [15–19], together with the ligand-binding domain (LBD) of Esrrb [20]. Recent studies depicted even the splendid structures of Oct4 and Sox2 DBDs binding to nucleosomes, hence deciphering their “pioneer factor” abilities [21–29]. Another structure of Sox2 bound to the importin Imp*α*3 has also been published, showing how its two Nuclear Localization Sequences (NLSs) flanking the DBD are involved in Sox2 nuclear import [30]. The other segments of OSNE have been predicted to be intrinsically disordered regions of proteins (IDRs) [31], i.e. they should have no stable tertiary fold when isolated [32–36].

These IDRs appear to have important roles in binding partners involved in epigenetic reprogramming, chromatin reorganization and in recruiting transcription or repression machineries [5,37–41]. These functions are poorly understood, and, to the best of our knowledge, no experimental characterization of the structural behavior of these regions in N- and C-terminal of DBDs has been released yet. Recent studies have shown that C-terminal regions of Oct4 and Sox2 are important for their reprogramming capacities [42, 43], notably by contributing to the engagement in molecular phase-separated condensates with the Mediator complex [44]. More generally, the activating or repressive activities of IDRs of transcription factors (TFs) have been scarcely studied at the structural level: these segments are thought to contain hydrophobic patches flanked by acidic amino acids, which favors DNA-binding specificity, phase separation and low-specificity interactions, notably with the Mediator subunit Med15 [44–53]; more specific interactions have been described in some cases [45,54,55].

Post-translational modifications (PTMs) add a layer of complexity, by being often responsible of the regulation of IDRs’ interactions [32,33,56,57], and notably of TFs activity [58–61]. Post-translational modifications (PTMs) are classical carriers of cell signaling by regulating the stability and the interactions of proteins. An increasing number of PTMs have been described on OSNE IDRs in the recent years [5,37,38,62–72] [73–77], notably phosphorylation by Cyclin-dependent kinases (CDKs) [66,78–87] or by Mitogen-activated protein kinases (MAPKs) [83,88,89], or their complementary Ser/Thr O-GlcNAcylation by OGT [65,90–96]. In order to prompt future studies on this topic, we thought to establish and test an experimental strategy i) for producing well-characterized samples of post-translationally modified IDRs of PluriTFs and ii) to use these as baits in pull-down assays for identifying PTMs’ related binding partners.

Hence, we characterized some of the phosphorylation reactions of Esrrb and Sox2 by p38*α*/*β*, Erk2 and Cdk1/2. Then, we showed that biotinylated chimera of Sox2 and Esrrb coupled to an AviTag peptide could be attached to streptavidin-coated beads. Finally, we loaded truncated segments of the C-terminal IDR of Sox2 (phosphorylated or not) on these beads, and exposed them to extracts of mouse Embryonic Stem Cells (mESCs) in pull-down assays, which we analyzed using quantitative mass spectrometry-based proteomics. Among the quantified (phospho)-Sox2 binders, we verified the phospho-dependent interaction between the proline cis-trans isomerase Pin1 and Sox2 using NMR spectroscopy.

## 2. Material and methods

### 2.1. Production of recombinant fragments of Oct4, Sox2, Nanog and Esrrb

We used human protein sequences, unless specified. Codon-optimized (for expressing in *Escherichia coli*) genes coding for human Oct4(aa1-145) and Oct4(aa286-380) were synthesized in the context of larger genes coding for Tev-Oct4(aa1-145)-Tev-GB1 and Tev-Oct4(aa286-380)-Tev-GB1 by Genscript and cloned into pET-41a(+) vector between SacII and HindIII restriction sites, hence permitting the expression of GST-His6-Tev1-Oct4(aa1-145)-Tev2-GB1 and GST-His6-Tev1-Oct4(aa286-380)-Tev2-GB1; Tev1 and Tev2 are the heptapeptide ENLYFQG cleavage site of the TEV protease, Tev2 is separated by GAGGAGG from GB1 (T2Q variant of the immunoglobulin binding domain B1 of the protein G from group G Streptococcus [97, 98]). The C-terminal GB1 tag was added to avoid any C-terminal proteolysis of the IDR of interest during the expression and the first purification steps; we did not test constructs without this supplementary folded domain, whose necessity for the stability of the IDR is thus not proven.

The same rationale (cDNA synthesis, cloning, vectors, chimera constructs) was used for producing Nanog(aa154-305), Nanog(aa154-215), Nanog(aa154-272), Nanog(aa154-305_ C185A-C227A-C243A-C251A), Nanog(aa154-272_C185A-C227A-C243A-C251A), and a very similar rationale (chimera constructs missing the C-terminal Tev2-GB1) for Sox2(aa1-42), Sox2(aa115-317_C265A), Sox2(aa115-187), Sox2(aa115-236), Sox2(aa115-282_C265A), Esrrb(aa1-102_C12A-C72A-C91A), Esrrb(aa1-102_C12A-C91A), Nanog(aa1-85) (this latter was cloned in the MfeI/HindIII restriction sites from pET-41a(+)).

The recombinant production and the purification of the protein constructs followed the procedures described previously [99], using the soluble fraction of bacterial lysates, except for the constructs containing the Sox2 C-terminal fragments. These latter constructs were recovered from the insoluble fractions of the lysates, and resolubilized in 8 M urea; these were submitted to a His-Trap purification in urea, and the last size-exclusion chromatography (SEC) had to be carried out in 2 M urea, which avoided clogging of the column and permitted to obtain regular elution peak widths (these were otherwise extremely broad, up to 100 mL for the longest Sox2(aa115-317_C265A) construct). The samples were concentrated and stored at −20 °C, and thawed just before the NMR experiments. The Sox2 samples containing 2 M urea were submitted to 2-3 cycles of concentration/dilution in Hepes at 20 mM, NaCl at 75 mM to generate samples in urea at 0.25 or 0.125 M.

All purification steps were carefully carried out at 4 °C; protein eluates from every purification step were immediately supplemented with protease inhibitors (EDTA-free cOmplete, Roche) (together with DTT at 10 mM for cysteine-containing protein constructs), before being submitted to a concentration preparing the next purification step. Chimera constructs of Sox2 and Esrrb IDR fragments containing a 15-mer peptide AviTag GLNDIFEAQKIEWHE were produced using procedures similar to those described earlier for OSNE constructs. The construct Sox2(aa234-317)-AviTag-His6 was soluble, and did not require to be purified in urea. More details about the production of OSNE peptides are given in the Supplementary Material.

### 2.2. Production of the biotin ligase BirA and specific biotinylation of the AviTag-peptide chimera

The biotin ligase BirA was produced using recombinant production in *E. coli* BL21(DE3)Star transformed with a pET21-a(+) plasmid containing a gene coding for BirA cloned at EcoRI and HindIII restriction sites. pET21a-BirA was a gift from Alice Ting (Addgene plasmid # 20857)[100]. The expression was carried out over night at 20 °C in a Luria-Bertani culture medium. The construct was containing a His6 tag in C-terminal and was purified using a two-step purification procedure including a His-trap followed by a SEC. Details about the production of BirA are given in the Supplementary Material.

The biotinylation was executed using a rationale inspired from a published protocol [101], at room temperature during 90 minutes, in samples containing the AviTag-chimera of interest at 100 μM and BirA at 0.7 μM in a buffer containing ATP at 2 mM, biotin at 600 μM, MgCl_2_ at 5 mM, DTT at 1 mM, HEPES at 50 mM, NaCl at 150 mM, protease inhibitors (EDTA-free cOmplete, Roche), at pH 7.0. To remove some eventual proteolyzed peptides and BirA, the biotinylated constructs were purified using a SEC in a column (Superdex 16/60 75 pg, Cytiva) preequilibrated with a buffer containing phosphate at 20 mM, NaCl at 150 mM at pH=7.4 (buffer called thereafter Phosphate Buffer Saline, PBS). The eluted fractions of interest were concentrated and stored at −20 °C.

### 2.3. Assignment of NMR signals from OSNE fragments, and structural propensities

The assignment strategy was the same than in previous reports from our laboratory [99]. The ^15^N relaxation data were recorded and analyzed according to the methods described in previous reports [102]. Details are given in the Supplementary Material.

Disorder prediction were calculated using the ODINPred website (https://st-protein.chem.au.dk/odinpred) [103]. Experimental secondary structure propensities of unmodified OSNE peptides were obtained using the neighbor-corrected structural propensity calculator ncSCP [104, 105] (http://www.protein-nmr.org/, https://st-protein02.chem.au.dk/ncSPC/) from the experimentally determined, DSS referenced C*α*and C*β* chemical shifts as input, with a correction for Gly-Pro motifs (−0.77 ppm instead of −2.0 ppm) [106]. Some signals were too weak in 3D-spectra from Sox2(aa115-317_C265A) recorded at 950 MHz, and their chemical shifts were not defined. In these cases, chemical shifts from 3D-spectra of Sox2(115-236) or His6-AviTag-Sox2(234-317_C265A) were used to complete the lists of chemical shifts used to calculate the chemical shift propensities shown in Figure 2.

### 2.4. NMR monitoring of phosphorylation reactions and production of phosphorylated peptides

We performed the phosphorylation kinetics presented in Figure 4a using commercial recombinant kinases GST-p38*β*at 10 μg/mL (Sigma-Aldrich, ref. B4437), GST-Erk2 at 20 μg/mL (Sigma-Aldrich, ref. E1283), GST-Cdk1/CyclinA2 at 20 μg/mL (Sigma-Aldrich, stock ref. C0244) and GST-Cdk2/CyclinA2 at 20 μg/mL (Sigma-Aldrich, ref. C0495). Then, we used kinases produced in house in *E. coli*, using plasmids containing optimized genes coding for p38*α*(aa1-360, full-length) and Erk2(aa8-360); these were produced, activated and purified in house as described previously [99]; in-house p38*α* was used at 40 μg/mL for the experiments shown in Figure 4.

Phosphorylation reactions were carried out using ^15^N-labeled IDRs at 50 μM, in Hepes 20 mM, NaCl 50 mM, DTT or TCEP at 4 mM, ATP 1.5 mM, MgCl_2_ at 5 mM, protease inhibitors (Roche), 7.5% D_2_O, pH6.8 at 25°C in 100 μL using 3 mm diameter Shigemi tubes. We monitored the phosphorylation kinetics by recording time series of ^1^H-^15^N SOFAST-HMQC spectra at 600 or 700 MHz, and quantifying the NMR signal intensities of the disappearing unphospho- and the appearing phospho-residues. We applied the methods that we described progressively in earlier publications [107–110]. More details are given in the Supplementary Material.

### 2.5. Pull-down assays

The extracts from mouse Embryonic Stem Cells (mESCs) were obtained from mESCs cultured in the conditions previously described [111]. Homogeneous extracts were obtained using DNA shearing by sonication in presence of benzonase, as described by Gingras and colleagues [112].

The pull-down assays were executed using 25 μL of streptavidin-coated magnetic beads (Magbeads streptavidine, Genscript) loaded with 1 nmol of the biotinylated bait-peptides of interest. These were incubated during one hour at room temperature with mESCs extracts, washed in PBS and eluted using a 2x Laemmli buffer. Details are given in the Supplementary Material.

### 2.6. Mass spectrometry-based proteomics analysis of pull-down assays

The pull-down samples were treated on-beads by trypsin/LysC (Promega). The resulting peptides were loaded and separated on a C18 columns for online liquid chromatography performed with an RSLCnano system (Ultimate 3000, Thermo Scientific) coupled to an Orbitrap Fusion Tribrid mass spectrometer (Thermo Scientific). Maximum allowed mass deviation was set to 10 ppm for monoisotopic precursor ions and 0.6 Da for MS/MS peaks. The resulting files were further processed using myProMS v3.9.3 (https://github.com/bioinfo-pf-curie/myproms; Poullet et al., 2007 [2]). False-discovery rate (FDR) was calculated using Percolator [3] and was set to 1% at the peptide level for the whole study. Label-free quantification was performed using peptide extracted ion chromatograms (XICs), computed with MassChroQ [4] v.2.2.1. The complete details are given in the Supplementary Material.

### 2.7. Recombinant production of Pin1 and NMR analysis of its interaction with Sox2 or phospho-Sox2

The plasmid containing the gene coding for the Pin1-WW domain was a kind gift from Isabelle Landrieu. The production was executed according to the previously published protocol [113]. The NMR analysis of binding with phosphoSox2(aa115-240) were performed at 283K and pH 7.0 with the GST-Pin1-WW construct and ^15^N-labeled Sox2(aa115-240) mixed in stoichiometric proportions, either at 50 or 10 μM for non-phospho and phosphoSox2, respectively. The details on the NMR acquisition, processing and analysis are given in the Supplementary Information.

## 3. Results

### 3.1. Structural characterization of the N- and C-terminal regions of Oct4

We produced and purified protein constructs containing the fragments of human Oct4(aa1-145) and Oct4(aa286-360), which were both predicted to be mostly disordered (Supp. Mat 3.2.3.). The 2D ^1^H-^15^N HSQC NMR spectra showed crosspeaks in the spectra region where random coil peptides resonances are usually found (Figure 1). We assigned the backbone NMR signals of ^1^H_N_, ^15^N, ^13^C*α*, ^13^C*β*, ^13^CO for both segments, which permitted to calculate experimentally derived secondary structure propensities from the ^13^C*α* and ^13^C*β* chemical shifts. We did not find any sign of a stable secondary structure, the highest *α*-helical propensities reaching about 25 % in short stretches of about 5 consecutive amino acids. Hence, we verified experimentally that these N- and C-terminal fragments of human Oct4 are IDRs.

**Figure 1.**
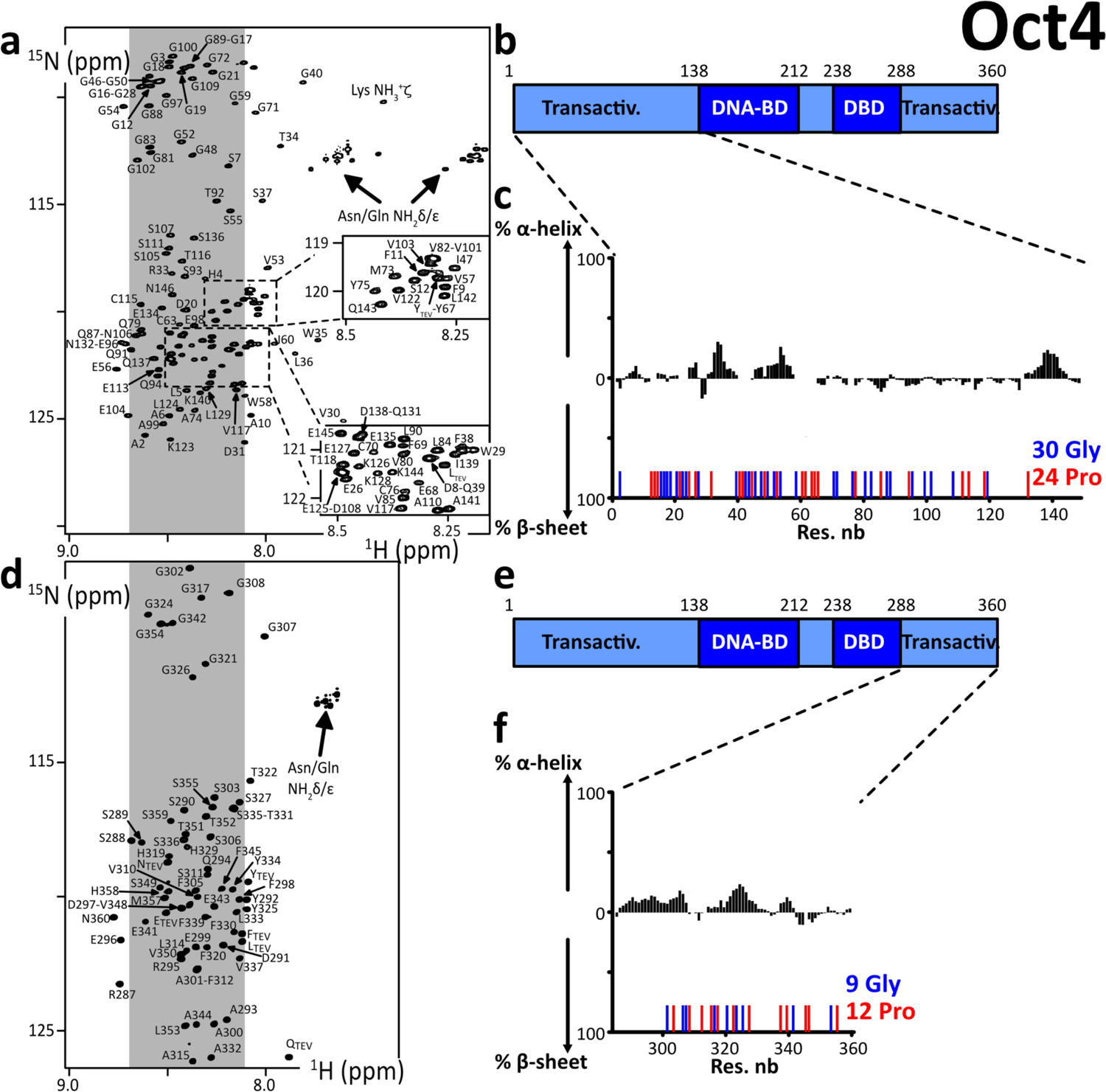
**a. d.** 2D ^1^H-^15^N HSQC spectra of the N- and C-terminal IDRs of human Oct4, the labels indicating the assignments; the grey areas show the spectral regions where random coil amino acids resonate usually; **b. c.** Primary structures of Oct4; dark and light colors indicate the folded and disordered domains, respectively; blue and red sticks indicate the positions of Glycines and Prolines, respectively; **c. e.** Secondary structure propensities calculated from the experimental chemical shifts of the peptide backbone Cα and Cβ, using the ncSPC algorithm [104, 105].

**Figure 2:**
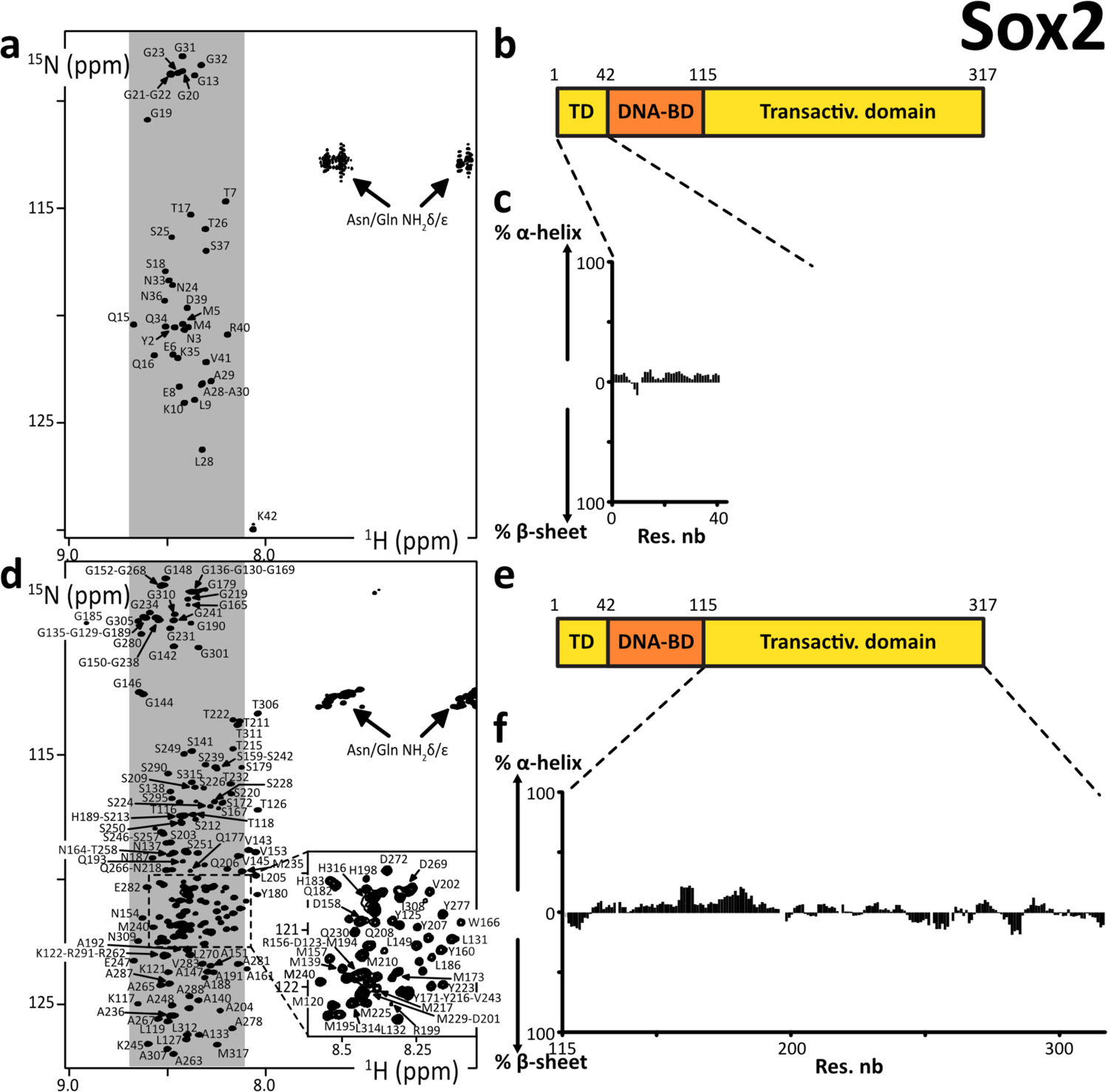
**a. d.** 2D ^1^H-^15^N HSQC spectra of the N- and C-terminal IDRs of human Sox2, the labels indicating the assignments; the grey areas show the spectral regions where random coil amino acids resonate usually; **b. c.** Primary structures of human Sox2; dark and light colors indicate the folded and disordered domains, respectively; **c. e.** Secondary structure propensities calculated from the experimental chemical shifts of the peptide backbone Cα and Cβ, using the ncSPC algorithm [104, 105].

We can notice that Oct4 IDRs contain a high density of Prolines, which are not directly detectable in the present ^1^H_N_-detected experiments, even though most of the ^13^C*α*, ^13^C*β* and resonances were characterized via HNCAB and HNCO experiments. We have shown previously that the ^13^C-detected experiments ^13^C*α*^13^CO permitted to observe all these Pro residues in Oct4(aa1-145) [99], whose chemical shifts were those of random coil peptides.

### 3.2. Structural characterization of the N- and C-terminal regions of Sox2

We produced and purified peptide fragments of human Sox2, namely Sox2(aa1-42), Sox2(aa115-187), Sox2(aa115-236), Sox2(aa115-282), Sox2(aa234-317_C265A), and Sox2(aa115-317_C265A). We also produced and purified chimera peptides His6-AviTag-Sox2(aa115-240) and His6-AviTag-Sox2(aa234-317_C265A). These were all predicted to be disordered (Supp. Mat 3.3.3.).

We had solubility issues with all of them but Sox2(aa1-42), Sox2(aa234-317_C265A) and His6-AviTag-Sox2(aa234-317_C265A). We had to recover these troublesome peptides from the insoluble fraction of the bacterial extract, after overexpression at 37 °C. We had even to carry out our final size-exclusion chromatography (SEC) purification step in a buffer containing urea at 2 M (at 4 °C) for Sox2(aa115-236), Sox2(aa115-282), His6-AviTag-Sox2(aa115-240), and Sox2(aa115-317-C265A). The assignments of these latter constructs were achieved in 0.25-0.5 M urea, after executing 2 to 3 concentration-dilution steps.

Aggregates were forming during the acquisition, which made the assignment rather painful. This behavior correlated with liquid-liquid phase separation (LLPS) propensities, which we observed a few months before such a behavior was reported by Young and collaborators [44]. The assignment of Sox2(aa115-317-C265A) was possible only at 950 MHz with the help of the previously assigned smaller fragments Sox2(aa115-236) and His6-AviTag-Sox2(aa234-317_C265A). Some stretches of amino acids were particularly difficult to observe in 3D spectra, e.g. the region aa160-185, because of an apparent fast T2 relaxation. We may investigate these phenomena in later reports.

We observed crosspeaks in the 2D ^1^H-^15^N HSQC NMR spectra that were all resonating in the spectral region of random coil peptides resonances (Figure 2). The spectra of the short fragments of the C-terminal region of Sox2 were exactly overlapping with those of Sox2(aa115-317_C265A) (Supp. Fig. 1). This shows that all these fragments have very similar local conformational behaviors, a phenomenon regularly observed with IDRs. We assigned the backbone NMR signals of ^1^H_N_, ^15^N, ^13^C*α*, ^13^C*β*, ^13^CO for Sox2(aa1-42), Sox2(aa115-236), His6-AviTag-Sox2(aa234-317_C265A) and partially for Sox2(aa115-317_C265A). We aggregated the lists of chemical shifts of the C-terminal fragments, and used them to calculate the experimental secondary structure propensities from the ^13^C*α* and ^13^C*β* chemical shifts. This confirmed the absence of any stable secondary structure elements in Sox2 N- and C-terminal region. The C-terminal region is poorly soluble below 0.25 M urea; this should not affect a stable fold, so we can affirm that these regions of Sox2 are experimentally proven IDRs.

### 3.3. Structural characterization of the N-terminal region of Nanog

We produced and purified the N-terminal peptide fragment of human Nanog(aa1-85). All our attempts to purify C-terminal regions of Nanog failed, even after alanine-mutation of cysteines in Nanog(aa154-305), Nanog(aa154-272) and Nanog(aa154-215). We managed to resolubilize our construct GST-His6-Tev-Nanog(aa154-305) from the insoluble fraction of the bacterial extract, to partially purify it and cleave it using the TEV protease. However, the resulting Nanog(aa154-305) peptide was barely soluble in a detergent (NP-40 at 2% v/v), and not in high salt buffers, or not even in presence of urea at 4 M. The 10 Tryptophane residues are probably playing a role in this behavior, in the context of a primary structure containing not enough hydrophobic amino acids favoring stable folds.

The crosspeaks of Nanog(aa1-85) in the 2D ^1^H-^15^N HSQC NMR spectrum were all in the spectral region of random coil peptides resonances (Figure 3). The assignment of the backbone NMR signals of ^1^H_N_, ^15^N, ^13^C*α*, ^13^C*β*, ^13^CO permitted to calculate experimental secondary structure propensities, which were low through the whole peptide. This Nanog N-terminal is thus confirmed to be an IDR.

**Figure 3:**
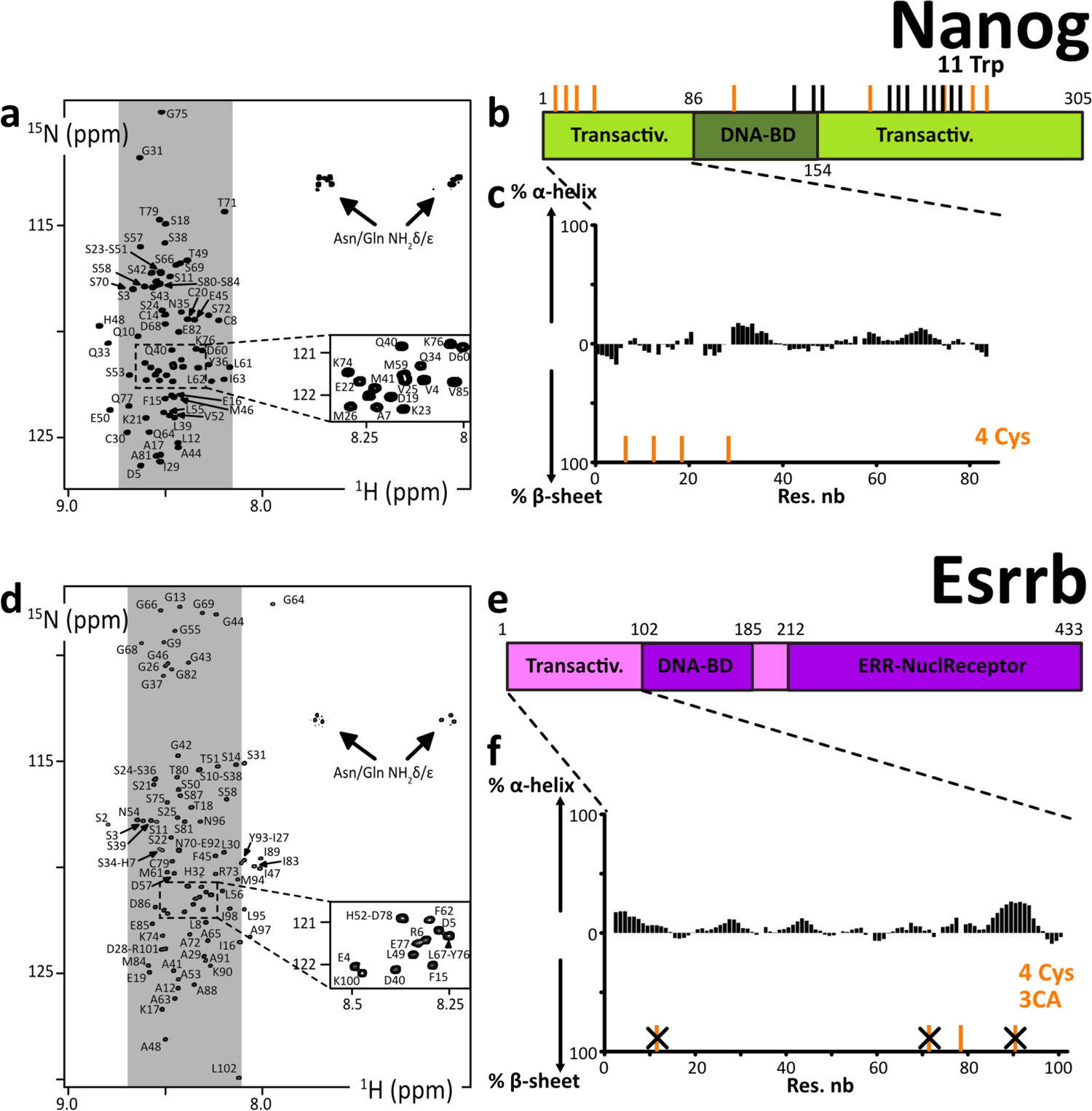
**a. d.** 2D ^1^H-^15^N HSQC spectra of the N-terminal IDRs of human Nanog and Esrrb, the labels indicating the assignments; the grey areas show the spectral regions where random coil amino acids resonate usually; **b. c.** Primary structures of human Nanog and Esrrb; dark and light colors indicate the folded and disordered domains, respectively; orange and black sticks indicate the positions of cysteines and tryptophanes, respectively; **c. e.** Secondary structure propensities calculated from the experimental chemical shifts of the peptide backbone Cα and Cβ, using the ncSPC algorithm [104, 105].

### 3.4. Structural characterization of the N-terminal region of Esrrb

We produced and purified the N-terminal fragment of human Esrrb(aa1-102), which was predicted to be disordered (Supp Mat. 3.5.3.), in the alanine-mutated versions Esrrb(aa1-102_C12A-C72A-C92A), Esrrb(aa1-102_C12A-C92A). This was a strategic choice to attenuate the formation of disulfide bonds; the wild-type N-terminal fragments might however be workable too. Mutating cysteines permitted to work in more comfortable conditions and to maintain our construct monomeric for longer periods of time in the next phosphorylation and biotinylation experiments. Cysteines are indeed highly solvent-accessible in IDRs and they are consequently difficult to keep in their thiol, non-disulfide forms, even in presence of fresh DTT or TCEP at neutral pH. We also produced chimera constructs Esrrb(aa1-102_C12A-C72A-C92A)-AviTag-His6 and Esrrb(aa1-102_C12A-C72A)-AviTag-His6.

All the 2D ^1^H-^15^N HSQC NMR spectra revealed crosspeaks in the spectral region of random coil peptides (Figure 3). These spectra are overlapping to a large extent, confirming the weak influence of the mutations of cysteines: the mutation Cys72Ala has almost no consequences on the chemical shifts, below 0.05 ppm even for the neighboring amino acids (Supp. Fig 2a,b); the mutation Cys91Ala has more impact, with chemical shifts perturbations of about 0.1 ppm for the next 5 amino acids (Supp. Fig. 2c,d), which is at least partially due to the fact that the Ala substitution favors an increase of local *α*-helicity (about 25 %, see Supp. Fig. 2e,f). This N-terminal fragment of human Esrrb is thus an IDR, according to the calculated secondary structure propensities (Figure 3f, Supp. Fig. 2e,f).

### 3.5. Phosphorylation of Esrrb and Sox2 by p38α, Erk2, Cdk1/2 as monitored by NMR spectroscopy

We reported recently the site-specific phosphorylation kinetics of Oct4 by p38*α* using ^13^C-direct NMR detection [99]. Here, we used more standard ^1^H-detected/^15^N-filtered experiments to rapidly characterize the site preferences of MAPKs and CDKs on Esrrb and Sox2, which we thought to use as baits for performing phospho-dependent pull-down assays (see below).

To start with, we used commercial aliquots of MAPKs, namely p38*β* and Erk2, and CDKs, namely Cdk1/CyclinA2 and Cdk2/CyclinA2, on Esrrb(aa1-102). We observed the progressive phosphorylation of its three Ser-Pro motifs, i.e. at Ser22, Ser34 and Ser58, in agreement with the consensus motifs of these kinases [114, 115]. Ser22 is the preferred target in all cases, while Ser34 is the least processed by CDKs, if at all: the commercial CDKs are poorly active in our hands, which we have verified with a number of other targets for years in the laboratory; this makes it difficult to distinguish between sites that are only mildly disfavored or those that are more stringently ignored by CDKs in NMR monitored assays.

The limited activities and high costs of commercial kinases motivated us to develop in-house capacities in kinase production. p38*α* was the most accessible to produce among the MAPKs and CDKs; we produced it and activated it using recombinant MKK6. We tested our home-made p38*α* on His6-AviTag-Sox2(aa115-240) and His6-AviTag-Sox2(aa234-317). It phosphorylated all the Ser/Thr-Pro motifs of these two peptides, and also T306 in a PGT_306_AI context, which shows a favorable Proline in position −2 [116], and a less common S212 in a MTS_212_SQ context.

Hence, we were able to generate AviTag-IDR chimera in well-defined phosphorylated states. To produce phosphorylated ^14^N-AviTag-IDR dedicated to pull-down assays, we executed the same protocol, and monitored in parallel “identical”, but ^15^N-labeled samples by NMR. Hence, we could no the exact phosphorylation status of the ^14^N-AviTag-IDR for the next experiments.

### 3.6. Structural characterization of AviTag-peptide chimera and biotinylated versions thereof

We aimed at identifying new partners of OSNE using pull-down assays. We thought to use chimera containing GST at the N-terminus, which appeared as a convenient approach: vectors integrating a GST-coding DNA sequence for overexpression in *E. coli* are available and of common use; glutathione-coated beads are also accessible and permit efficient and specific binding of GST-containing chimera peptides. However, we were unsatisfied by the performances of the method: GST binding to glutathione-coated beads is slow at low temperature (necessary to avoid IDR proteolysis); moreover, it appeared that GST-IDRs chimera are hampered by the IDR “molecular-cloud” and are even weaker and slower to bind the beads. Our attempts to bind GST-IDRs to the beads were thus resulting in poor yields, which were not very reproducible. In the context of our aims, i.e. to establish a method allowing semi-quantitative detection of IDR binding partners, this unsatisfying lack of reproducibility was only promising supplementary variable parameters.

Thus, we decided to switch to another strategy: the use of the specifically biotinylated 15-mer peptide tag called AviTag [101]. This is efficiently and specifically biotinylated by the biotin ligase BirA (Figure 5d), which permits the high-affinity binding to streptavidin-coated beads. We designed AviTag-IDR chimera, with the AviTag in N-terminal position for Sox2-IDRs constructs, and in C-terminal for Essrb-IDR constructs. We characterized the AviTag and its impact on the IDRs of interest using NMR: the AviTag is unfolded and it does not affect the Sox2 and Esrrb fragments, according to the observed negligible chemical shift perturbations (Fig 5) – the GST-Tag provoked also only very weak chemical shift changes on Essrb(aa1-102) (Fig S3). The biotinylated AviTag peptide appears to be slightly less mobile than the common IDPs on the ps-ns timescale, according to the heteronuclear ^1^H^15^N-nOes (Figure 5b). We observed that the biotinylation of the AviTag provokes weak, but distinguishable chemical shift perturbations for the close neighbors of the biotinylated lysine, but had no effect on the peptides of interest (Figure 5c). It generated also the appearance of a HN-ester NMR signal, similar to that of acetylated lysine [107, 117]. Hence, we could quantify and monitor the reaction advancement using NMR, and determine the incubation time that was necessary and sufficient to obtain a complete biotinylation of our chimera AviTag-IDRs (see 2.2.). This was one among many optimization steps permitting the production of sufficient quantities of intact IDRs for the pull-down assays.

**Figure 4:**
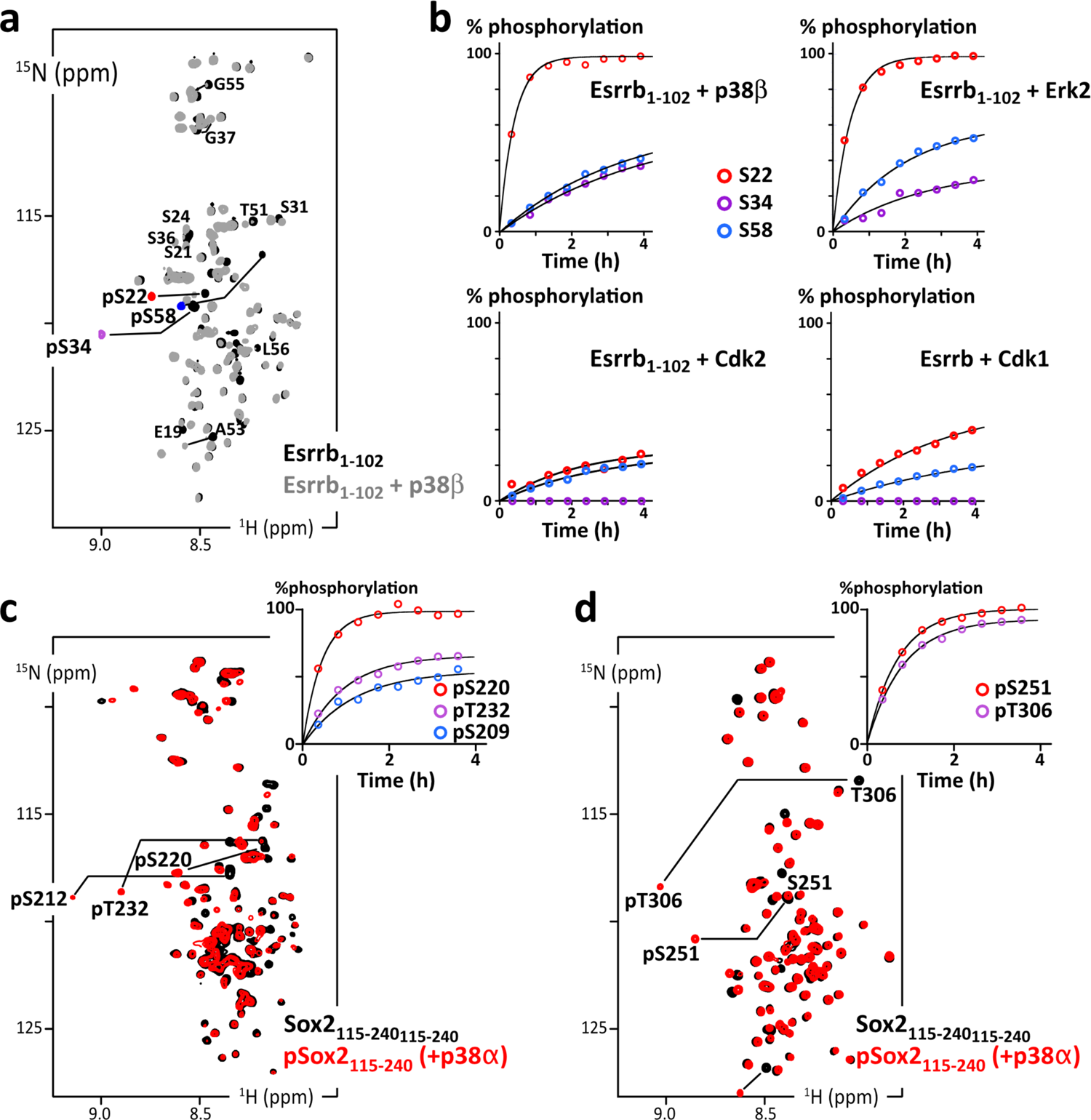
**a.** Overlay of 2D ^1^H-^15^N HSQC spectra of Esrrb(aa1-102_C12A-C72A-C91A) before (black) and after (grey and phosphosites colored in red/purple/blue) phosphorylation by p38β; **b.** Residue specific time courses of the phosphorylation of Esrrb(aa1-102) executed by commercial kinases p38β, Erk2, Cdk2 or Cdk1, as measured in time series of 2D ^1^H-^15^N SOFAST-HMQC spectra recorded during the reaction; **c.** Overlay of 2D ^1^H-^15^N HSQC spectra of AviTag-Sox2(aa115-240) before (black) and after (red) phosphorylation by p38a; the inset at the top-right shows the residue specific phosphorylation build-up curves, as measured in time series of 2D ^1^H-^15^N SOFAST-HMQC spectra; d. Same as **c.** with AviTag-Sox2(aa234-317_C265A).

**Figure 5.**
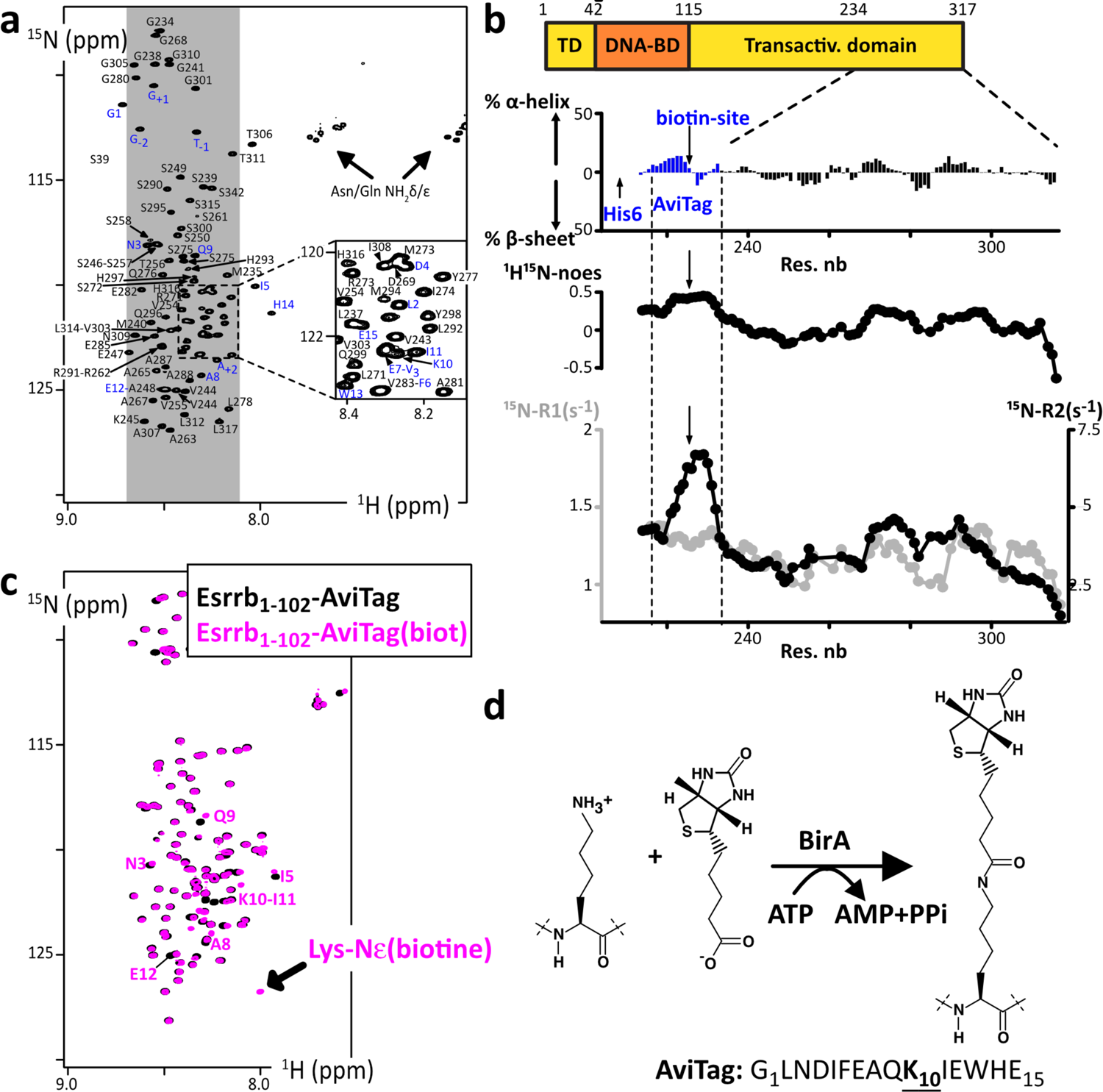
**a.** 2D ^1^H-^15^N HSQC spectrum of ^15^N-His6-AviTag-Sox2(aa234-317_C265A), the blue labels indicated the assigned signals from the AviTag residues; **b.** Secondary structure propensities calculated from the experimental chemical shifts of the peptide backbone Cα and Cβ, using the ncSPC algorithm [104, 105]; the residue specific ^1^H-^15^N-noes, ^15^N-R1 (grey) and ^15^NR2 (black) measured at 600 MHz are shown below (the profiles show values averaged over three consecutive residues); **c.** Overlay of 2D ^1^H-^15^N HSQC spectra of the Esrrb(aa1-102_C12A-C72A-C91A)-AviTag-His6 before (black) et after (magenta) biotinylation by BirA; the NMR signals from the residues neighboring the biotinylation site are indicated, which permit the quantification of the biotinylated population; **d.** Scheme of the reaction of Avi-Tag biotinylation executed by the ATP-dependent BirA.

Next, we tested the binding of the biotinylated AviTag-IDRs on streptavidin-coated beads. This produced very satisfying results, i.e. stoichiometric, specific binding in one hour with no leakage (Fig. S4). This approach was thus selected for the pull-down assays.

### 3.7. Identification of Sox2 binding partners in extracts from mouse Embryonic Stem Cells

We prepared the four peptides AviTag-Sox2(aa115-240) and AviTag-Sox2(aa234-317) in their non-phosphorylated and phosphorylated versions, using p38*α* to execute the phosphorylation reactions (Figure 6a). These peptides were also biotinylated, and later bound to streptavidin-coated beads, which we used as baits for pull-down assays in extracts from mESCs (Figure 6b). Importantly, size-exclusion chromatographies were carried out between every step to discard proteolyzed peptides, the enzymes (kinases of BirA) and their contaminants as much as possible. We performed the pull-down assays with the four samples in parallel with the same cell extract, in duplicate, and then analyzed the bound fractions using quantitative LC-MS/MS analysis (see Supp. Mat 1.6. for the full description). Hence, we could identify and evaluate the relative quantities of proteins retained by the four AviTag-peptides (Figure 6). On paper, this presents the important advantage of removing false-positive binders, which can interact unspecifically with the streptavidin-coated beads.

**Figure 6:**
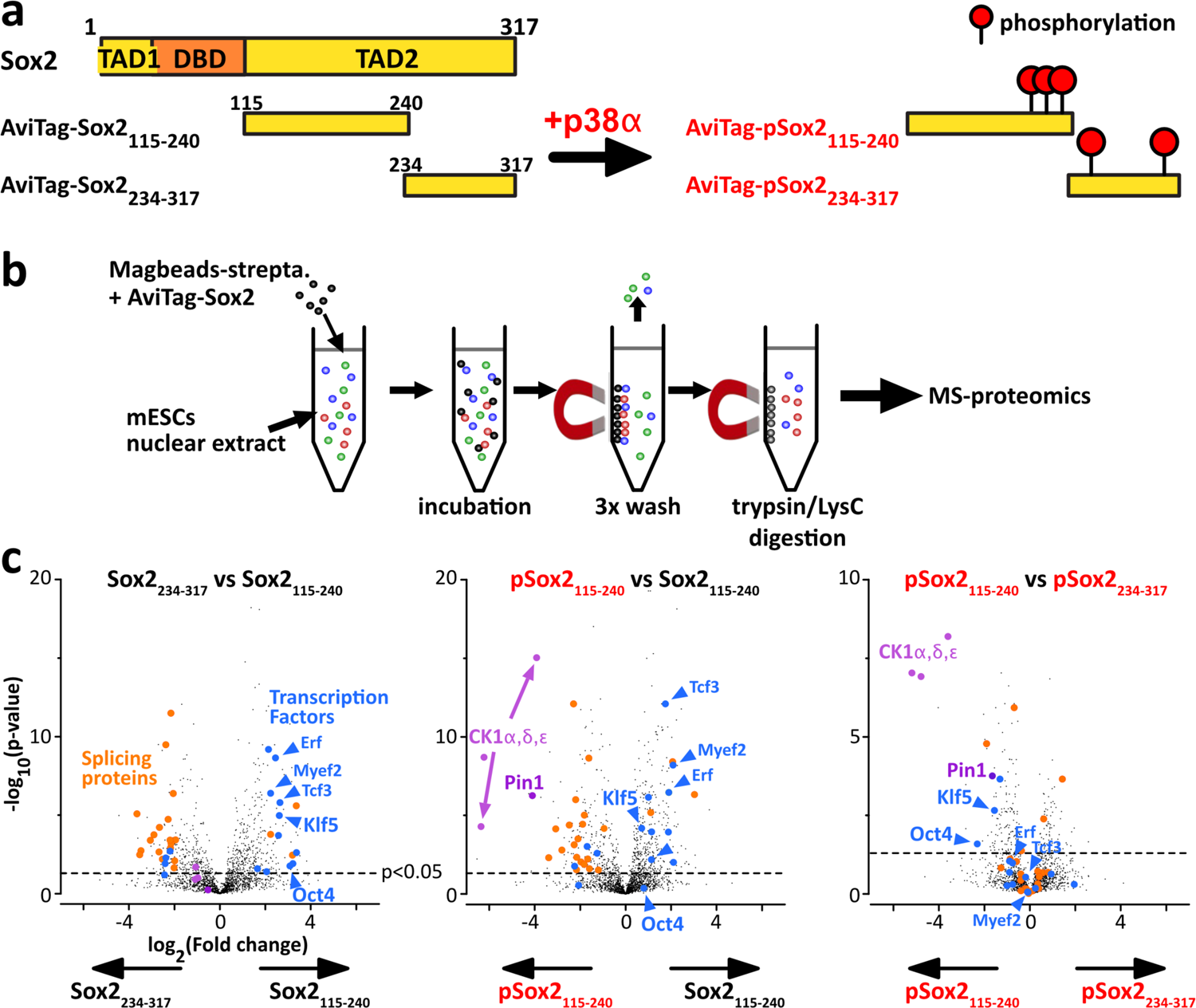
Differential interactomics of Sox2 constructs upon p38α phosphorylation. **a.** We have produced unmodified/phospho-Sox2 truncations carrying N-ter biotinylation on the AviTag, and later attached on streptavidin-magnetic beads. **b.** We have generated mESCs nuclear extracts for pull-down assays using our Sox2 constructs as baits. **c.** The volcano plots of the log2 ratios, showing a quantitative analysis of the proteins present at the end of the pull-down assays; the dashed line indicates the threshold of p-value<0.05; we highlighted interesting partners in: <u>blue:</u> transcription factors associating with Sox2(aa115-240); <u>magenta:</u> phospho-dependent partners of Sox2(aa115-240) and or Sox2(aa234-317). Pull-downs have been performed in duplicates, using 15 million cells per sample (extract protein conc.: 5 mg/mL), and 1 nmol of bait protein. Experimental conditions may be improved (higher number of replicates, cells, washing conditions, …).

We present here results that should be interpreted carefully: we produced only duplicates for every condition, using one single cell extract. To deliver trustful information, the common standards in the field recommend 3 to 5 replicates, and the use of independent samples. We can still comment the results, which we consider to be interesting preliminary data. We quantified proteins with at least 2 distinct peptides across the two replicates (Figure 6C). Among the relative quantifications with an adjusted p-value < 0.05, we observed a two-fold change or more (FC>2) of a number of transcription factors in the proteins pulled out by AviTag-Sox2(aa115-240), among which the PluriTFs Oct4 and Klf5 are significantly enriched (>4 peptides, p-value<0.02). Most of these TFs appear to bind slightly less to the phosphorylated pSox2(aa115-240). However, pSox2(aa115-240) kept apparently a capacity to bind Oct4 and Klf5 to some extent, in comparison to pSox2(aa234-317). Also, we found out that pSox2(aa115-240) was pulling out the three isoforms of CK1 (>3 peptides, p-values<10^-5^) and the proline isomerase Pin1 (>2 peptides, p-values<2.10^-4^), while Sox2(aa115-240) and pSox2(aa234-317) did not or much less.

### 3.8. NMR characterization of the interaction between Pin1 and phospho-Sox2

We decided to test the interaction between pSox2(aa115-240) and Pin1. We recorded ^1^H-^15^N NMR spectra of ^15^N-labeled Sox2(aa115-240) or pSox2(aa115-240) alone or in presence of the Pin1-WW domain (natural abundance peptide, i.e. 0.6% ^15^N, 99.4% ^14^N, hence “NMR invisible” in ^15^N-filtered experiments). We observed localized losses in signal intensities for the residues neighboring the three pSox2(aa115-240) phosphosites when mixed with Pin1 (Figure 7b); in contrast, no significant differences showed up in the spectra obtained with non-phosphorylated Sox2(aa115-240) in absence or presence of Pin1-WW (Figure 7a).

**Figure 7:**
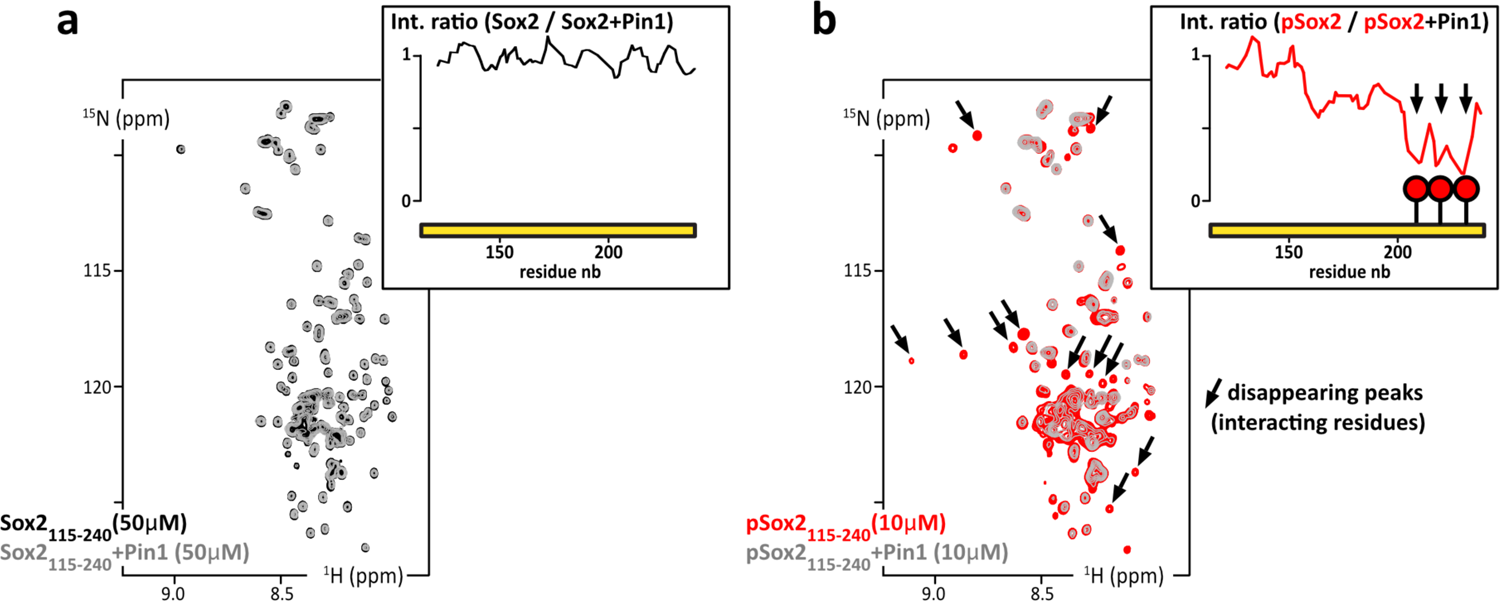
**a.** Overlay of 2D ^1^H-^15^N HSQC spectra of ^15^N-Sox2(aa115-240) alone at 50 μM (black) or mixed with Pin1 in isotopic natural abundance and in stoichiometric amounts (grey); inset up-right: residue specific NMR signal intensity ratios as measured in the two HSQC spectra. **b.** Overlay of 2D ^1^H-^15^N HSQC spectra of ^15^N-phosphoSox2(aa115-240) alone at 10 μM (red) or mixed with the Pin1-WW domain in stoichiometric amounts (grey); insets, up-right: residue specific NMR signal intensity ratios as measured in the two HSQC spectra (in absence/presence of Pin1-WW domain).

Hence, these signal losses are the typical signs of a position-specific interaction between an IDR and a folded protein in the intermediate or slow NMR time-scale, i.e. μs-s timescale, which corresponds to submicromolar affinities for this type of molecules. This shows that our pull-down assays were capable of isolating and identifying binding partners of an IDR of Sox2 in this range of affinities.

## 4. Discussion

The structural biochemistry analysis reported here can be applied to the broad field of transcription factors (TFs). These are essential actors of cell signaling: they are key elements for inducing or maintaining pluripotency or differentiation, for cell proliferation or cell-cycle arrest, by activating or repressing gene transcription [61, 118]. About 90% of the ∼1,600 TFs contain large disordered segments (> 30 consecutive amino acids), which is particularly true for PluriTFs [31, 119], and this correlation between TFs and IDRs exist in all kingdoms of life [45, 120]. IDRs of TFs recruit transcription co-factors or the transcription machinery, which is still not very well characterized in detail [44–55].

Indeed, the fine understanding of TFs interactions via their IDRs appears to be hampered by the nature of these interactions: weak affinities, multivalency, possible redundancy between TFs and coacervation; post-translational modifications (PTMs), which can switch on or off IDRs’ interactions, are a supplementary source of confusion when searching for binding partners. Among the difficulties, we should also mention the basic biochemistry issues: IDRs are difficult produce and manipulate *in vitro*, because they are prone to degrade or aggregate. Here, we have tried to demonstrate the feasibility and the interest of some biochemical and spectroscopic approaches to better characterize IDRs of TFs, their phosphorylation and the associated binding partners.

Like other TFs, pluripotency TFs Oct4, Sox2, Nanog and Esrrb are post-translationally modified (see the introduction), notably by CDKs and MAPKs [66,78–89]. These two classes of kinases are fundamental actors in all aspects of eukaryotic cellular life, and understanding their activity and regulation in pluripotency or differentiation is of high significance. Interesting questions are still pending: what is the phosphorylation status of OSNEs’ IDRs in pluripotent cells, what is the impact on their interaction networks, and how does it affect pluripotency or differentiation? The inhibition of MAPK Erk signaling is necessary to maintain pluripotency in the standard culture conditions of ESCs and iPSCs [66,82–84], while these cells show a high CDK activity [8,86,87]; these two kinases family have the same core consensus sites, i.e. Ser/Thr-Pro motifs, which are abundant in OSNEs’ IDRs and whose phosphorylation has apparently consequences for initiating differentiation [73,78,80,83,85]. We have shown that we could produce well-defined phosphorylation status of these peptides, using recombinant kinases and NMR analysis, which makes it possible to study their interactions *in vitro*. We have also demonstrated our capacity to use these peptides as baits in pull-down assays for detecting new binding partners.

Using this approach, we have identified an interaction between phospho-Sox2(aa115-240)-pS212-pS220-pT232 and the prolyl cis-trans isomerase Pin1. Interactions between Pin1 and Oct4 or Nanog have been published earlier [81, 121]. This does not represent a major surprise, because Pin1 recognize the consensus motif pSer/pThr-Pro, which is very degenerate (about one third of all the detected sites in phosphoproteomics) [122, 123]; dozens of Pin1 binding partners have been reported, which leaves suspicions about the functionality of all these, and about the capacity of the intracellular pool of Pin1 molecuels to engage in so many interactions. We also detected an affinity between phospho-Sox2(aa115-240)-pS212-pS220-pT232 and three CK1 isoforms. This capacity of CK1 kinases to recognize prephosphorylated in position −3 of a potential Ser/Thr is well-known (with a hydrophobic residue in +1), and a recent structural characterization of CK1*δ* has shown that it was even capable of keeping some affinity for a phosphorylated product after the phosphate transfer [124].

Hence, we feel that these results are at the same time satisfying, but do reveal degenerate interactions, whose biological significance is questionable. This corresponds to one of the major drawbacks in the field of IDRs’ studies: they generate interactions of weak affinities, which can be easily released during the washes of our pull-down assays. In this regard, the « proximity labeling » approaches (BioID, APEX and their derivates) appeared recently to be quite adapted to transient interactions: these methods, developed in the last ten years, use chimera constructs containing enzymes that transfer chemical groups to their intracellular neighbors, which can be later identified by mass-spectrometry [125–128]. IDRs are very flexible, solvent-exposed and establish a lot of poorly specific transient interactions. This was raising concerns about the possible production of many false-positive if one used proximity labeling methods to detect IDRs’ binding partners. This has been partially confirmed by a recent study, but this bias appears to be limited [129]. Interactomes of 109 TFs have actually been described using Bio-ID and affinity-purification MS, showing the complementarity between proximity-labeling and the pull-down approach proposed here [130]. Yeast double-hybrid, which can detect ∼20 μM affinity interactions, and novel phage display approaches will also help in this task [131–134].

Another difficulty in studying IDRs of TFs is their propensity to coacervate [44–53]. Here, we have tried to use Sox2 as a bait in pull-down assays, a protein that has been later recognized to favor liquid-liquid phase separation [44, 135]. We met this difficulty during the production steps, which forced us to purify most of the Sox2 constructs in urea at 2 M. We could straightforwardly observe liquid-liquid phase separation of Sox2(aa115-317) at 4 μM using DIC microscopy in presence of Ficoll-70 (data not shown), but also progressive aggregation and low solubility thresholds while working with our purified samples. These are clear limiting factors for sample production and NMR characterization, which will hamper a number of other studies on IDRs of TFs. This might also affect the results of pull-down assays: we noticed an enrichment in TFs in the samples obtained from pull-downs using Sox2(aa115-240) as a bait, which has a much higher coacervation propensity than Sox2(aa234-317). Is it possible to generate local surface liquid-liquid phase separation on the surface of streptavidin-coated beads? This might be at the same time a bless and a curse for future studies, by helping the formation of biologically significant assemblies, or by favoring unspecific, non-native macromolecules interactions.

A final bottleneck in the studies of these IDRs is the capacity to produce post-translationally modified samples. The commercial enzymes are not very well adapted to our NMR studies, because of the required quantities. Here, we have used in-house production of the kinase p38*α*. Since we carried out the present work, we have developed our capacities in producing activated Erk2, Cdk2/CyclinA1 and Cdk1/CyclinB1. These will be part of our future studies. TFs are indeed quite adapted to NMR investigations: they are 300 to 500 residues long and contain large IDRs (∼100 amino acids) separating small folded domains (also ∼100 amino acids) [31, 45]. Their structural characterization would permit to understand a number of cell signaling mechanisms at the atomic scale, and possibly to identify new therapeutic targets, even though this class of proteins is notoriously difficult to inhibit [60,136,137].

## 5. Conclusion

We have applied NMR techniques to carry out a primary analysis of the pluripotency transcription factors Oct4, Sox2, Nanog and Esrrb, in particular of their intrinsically disordered regions. We have shown experimentally that they did not adopt a stable fold when isolated, and that we were able to conduct a residue-specific analysis. This relies on the delicate production and purification of these peptides, which are prone to proteolysis and aggregation; producing them in a well-defined post-translational modification status was an even arduous challenge. We have demonstrated the feasibility of these tasks using recombinant kinases and NMR analysis. We have also evaluated the usefulness of such protein constructs as baits in pull-down assays to identify new binding partners of IDRs. These characterizations and the associated methods provide firm basis for future investigations on transcription factors.

## Declaration of Competing Interest

The authors declare that they have no competing interest.

## Fundings

This work was supported by the CNRS and the CEA-Saclay, by the French Infrastructure for Integrated Structural Biology (https://frisbi.eu/, grant number ANR-10-INSB-05-01, Acronym FRISBI) and by the French National Research Agency (ANR; research grants ANR-14-ACHN-0015 and ANR-20-CE92-0013). Financial support from the IR INFRANALYTICS FR2054 for conducting the research is also gratefully acknowledged. This work was also supported by grants from the “Région Ile-de-France” and Fondation pour la Recherche Médicale (D.L. and LSMP).

## Acknowledgements

We thank Thaleia Papadopoulou, Amandine Molliex and Navarro Pablo for providing mouse Embryonic Stem Cells (mESCs) extracts, and for fruitful discussions. We thank Nadia Izadi-Pruneyre for providing the bench space necessary to carry out the pull-down experiments with fresh mESCs extracts. We thank the IR-RMN (now Infranalytics), notably Nelly Morellet, François Giraud and Ewen Lescop, for their reactivity, their support, and their long-standing, efficient care of the 950 MHz spectrometer. We thank Marie Sorin, Baptiste Nguyen and Benjamin Bacri, who contributed to this study during their Master1-Master2 internships.

## Supplementary material

Supporting information for this article is available online at …

## Notes

### Competing Interest Statement

The authors have declared no competing interest.

