## Supplementary Material for "Structural characterization of stem cell factors Oct4, Sox2, Nanog and Esrrb disordered domains, and a method to identify their phospho-dependent binding partners"

### 1. Experimental procedures

#### 1.1. OSNE peptides production

The TEV (Tobacco Etch Virus) protease was produced in-house recombinantly in *E. coli* BL21(DE3)Star, from a construct containing a hexahistidine tag (His6).
All peptides were produced in *E. coli* (strain BL21(DE3)Star) transformed with the plasmids presented in the main text. Cells were grown in M9 medium containing $^{15}NH_4^+$ (0.5 g/L) and $^{13}$C-glucose (2 g/L) as sole sources of nitrogen and carbon for producing samples used for NMR assignment, and natural abundance $^{12}$C-glucose (2 g/L) otherwise. Media were supplemented with kanamycine at 50 µg/mL, and the expression was induced at an optical density $OD_{600}$=0.8 by supplementing the medium with ITPG at 1 mM at 37°C. Cells were harvested by centrifugation (5 minutes at 5,000 g) 4 hours later and cell pellets were stored at -20°C. Cells were lysed using sonication in Tris 20 mM, NaCl 150 mM, at pH 7.4 (buffer called "Tris Buffer Saline", TBS) in presence of benzonase (E1014 Sigma-Aldrich), lysozyme, protease inhibitors 1x (EDTA-free cOmplete, Roche) and 10 mM DTT.
Soluble and insoluble fractions were separated by 15 min of centrifugation at 15,000g. Oct4-, Nanog- and Esrrb-peptides were purified from the soluble fractions. The lysates were loaded on a His-Trap FF column (5 mL, Cytiva) and eluted using a gradient of imidazole (in TBS). The eluted fractions were concentrated, submitted to TEV treatment for 1 hour in TBS+imidazole supplemented with 10 mM DTT, and then diluted in TBS and re-loaded on the His-Trap column. Fractions containing the peptide of interest were submitted to a size-exclusion chromatography (SEC) in a column (Superdex 16/60 75 pg, Cytiva) previously equilibrated with Hepes at 10 mM or 20 mM, or phosphate at 20 mM, and NaCl 50 mM or 150 mM, at pH 6.8 (low-salt samples for NMR assignments, high-salt samples for phosphorylation kinetics or pull-down assays). For the cysteine-containing peptides, the eluted fractions of interest were immediately supplemented with DTT or TCEP at 2 mM, concentrated and stored at -20°C. Fresh DTT or TCEP was supplemented further after thawing before the NMR experiments.
The constructs containing Sox2(aa1-42), Sox2(aa115-187) and Sox2(aa234-317)-AviTag-His6 were also purified from the soluble fraction, as explained above. The other Sox2-constructs were recovered from the insoluble fractions of the lysates, and resolubilized in TBS supplemented with 8 M urea, loaded on a His-Trap FF column (5 mL, Cytiva) and eluted using a gradient of imidazole; the eluates were then supplemented with β-mercaptoethanol at 50 mM and incubated at room temperature for 15 minutes, before being dialyzed in TBS supplemented with DTT at 1 mM, in order to refold the GST domain. The samples were then submitted to TEV cleavage in 0.5 M urea, and a second His-Trap purification was carried out in TBS supplemented with urea at 2 M. The fractions of interest were concentrated and submitted to a SEC in Hepes at 10 mM or phosphate at 20 mM, and NaCl 50 mM or 150 mM, urea at 2 M, at pH 6.8. The samples were concentrated and stored at -20°C. Before the NMR experiments, they were thawed and submitted to 2-3 cycles of concentration/dilution in Hepes at 20 mM, NaCl at 75 mM to generate samples in urea at 0.25 or 0.125 M. We paid attention to avoid precipitation during the concentration steps, because these peptides had a limited solubility, about 100-150 µM.
We achieved some liquid-liquid phase separation assays, using DIC microscopy at room temperature in Ficoll-70 at 100 mg/mL. These were carried out with Sox2 peptides previously centrifuged during 10 minutes at 15,000g to remove the aggregates: for example, coacervates were observed at 4 µM of Sox2(aa115-317_C265A), and some aggregates were rapidly forming under the microscope at 20 µM.

#### 1.2. Production of BirA and biotinylation of AviTag-peptide chimera

Bacteria transformed with pET21a-BirA were precultured at 37 °C overnight in a Luria-Bertani (LB) culture medium supplemented with ampicillin at 100 µg/mL. Then, these were cultured in a larger volume of LB supplemented with ampicillin at 50 µg/mL at 37 °C, and at 30 °C when they reached an optical density (OD) of 0.4. At an OD=0.8, the culture was transferred to 20 °C and the protein expression was induced by supplementing the medium with IPTG at 0.5 mM. The incubation was carried out overnight, the bacteria were harvested by centrifugation at 4,500 g for 5 minutes and the pellets were stored at -20 °C.
The purification was carried out at 4 °C. Cells were lysed using sonication in TBS at pH 7.5 in presence of 0.5 µL of benzonase (E1014 Sigma-Aldrich), lysozyme, PMSF at 1 mM (Sigma-Aldrich) and DTT at 10 mM. The soluble and insoluble fractions were separated by 15 min centrifugation at 15,000g. The lysate (supernatant, soluble fraction) was loaded on a His-Trap column (His-Trap FF 5 mL, Cytiva) and eluted in TBS using a gradient of imidazole. The eluted fractions of interest were concentrated in presence of DTT at 10 mM, and later submitted to

a SEC in a column (Superdex 16/60 75 pg, Cytiva) previously equilibrated with TBS at pH 7.5, supplemented with 10% v/v glycerol. The fractions of interest were concentrated in presence of DTT at 2 mM. Final concentrations of BirA were about 100 µM. The obtained sample was aliquoted, flash-frozen and stored at -80 °C.
The primary sequence of the expressed construct is:

```
MKDNTVPLKLIALLANGEFHSGEQLGETLGMSRAAINKHIQTLRDWGVDVFTVPGKGYSLPEPIQLLNAKQILGQLDGGSVAVLPVIDSTNQYL
LDRIGELKSGDACIAEYQQAGRGRRGRKWFSPFGANLYLSMFWRLEQGPAAAIGLSLVIGIVMAEVLRKLGADKVRVKWPNDLYLQDRKLAGIL
VELTGKTGDAAQIVIGAGINMAMRRVEESVVNQGWITLQEAGINLDRNTLAAMLIRELRAALELFEQEGLAPYLSRWEKLDNFINRPVKLIIGD
KEIFGISRGIDKQGALLLEQDGIIKPWMGGEISLRSAEKKLAAALEHHHHHH*
```

#### 1.3. Assignment of NMR signals from OSNE fragments, and structural propensities

Almost all NMR spectra were recorded on a 700 MHz Bruker Avance Neo spectrometer or a 600 MHz Bruker Avance II, equipped with cryogenically cooled triple resonance $^{1}$H[$^{13}$C/$^{15}$N] probes optimized for $^{1}$H-detection, a TCI and a TXI, respectively. Assignment spectra of Sox2_aa115-317 were recorded on a 950 MHz Bruker Avance III spectrometer, equipped with a cryogenically cooled triple resonance $^{1}$H[$^{13}$C/$^{15}$N] probe (TCI). All spectra were processed in Topspin 3 or Topspin 4. 3D spectra analysis was carried out using CccpNmr 2.4.2. DSS at 100 µM and 7.5% $D_2O$ were added in all samples.
NMR assignments of backbone amide resonances of uniformly-labeled peptides ($^{13}$C/$^{15}$N) was achieved using BEST-HNCO, -HN(CA)CO, -HNCACB,[4] and (H)N(CA)NH 3D experiments, in HEPES at 10 mM, NaCl at 50 mM, DTT or TCEP at 2 to 5 mM, at pH 6.8 and 283K, and at peptide concentrations ranging from 150 to 900 µM in 5 mm diameter Shigemi tubes.
Assignments of Oct4_aa286-360, Sox2_aa1-42, Nanog_aa1-85, Esrrb_aa1-102 were carried out at 700 MHz; those of Oct4_aa1-145, Sox2_aa115-236, His6-AviTag-Sox2_aa234-317_C265A at 600 MHz; those of Sox2_aa115-317_C265A at 950 MHz.
Oct4_aa286-360: interscan delay: 0.5 s
B-HNCO and were carried out with 1024 ($^{1}$H) x 96 ($^{13}$C) x 80 ($^{15}$N) complex points and sweep widths of 12.98 ppm ($^{1}$H), 8 ppm ($^{13}$C) and 22 ppm ($^{15}$N),
B-HN(CA)CO were carried out with 1024 ($^{1}$H) x 96 ($^{13}$C) x 64 ($^{15}$N) complex points and sweep widths of 12.98 ppm ($^{1}$H), 8 ppm ($^{13}$C) and 22 ppm ($^{15}$N),
B-HNCACB 1024 ($^{1}$H) x 128 ($^{13}$C) x 64 ($^{15}$N) complex points and sweep widths of 12.98 ppm ($^{1}$H), 60 ppm ($^{13}$C) and 22 ppm ($^{15}$N),
B-(H)N(CA)NH with 1024 ($^{1}$H) x 64 ($^{15}$N) x 64 ($^{15}$N) complex points and sweep widths of 12.98 ppm ($^{1}$H), and 22 ppm ($^{15}$N).
Sox2_aa1-42: interscan delay: 0.5 s
B-HNCO and was carried out with 2048 ($^{1}$H) x 88 ($^{13}$C) x 88 ($^{15}$N) complex points and sweep widths of 12.98 ppm ($^{1}$H), 8 ppm ($^{13}$C) and 26 ppm ($^{15}$N),
HN(CA)CO was carried out with 2048 ($^{1}$H) x 72 ($^{13}$C) x 72 ($^{15}$N) complex points and sweep widths of 12.98 ppm ($^{1}$H), 8 ppm ($^{13}$C) and 26 ppm ($^{15}$N),
B-HNCACB 2048 ($^{1}$H) x 80 ($^{13}$C) x 80 ($^{15}$N) complex points and sweep widths of 12.98 ppm ($^{1}$H), 60 ppm ($^{13}$C) and 26 ppm ($^{15}$N),
B-(H)N(CA)NH with 2048 ($^{1}$H) x 64 ($^{15}$N) x 64 ($^{15}$N) complex points and sweep widths of 12.98 ppm ($^{1}$H), and 26 ppm ($^{15}$N).
Esrrb_aa1-102_3Cys->3Ala: interscan delay: 0.5 s
B-HNCO and was carried out with 2048 ($^{1}$H) x 92 ($^{13}$C) x 92 ($^{15}$N) complex points and sweep widths of 12.98 ppm ($^{1}$H), 8 ppm ($^{13}$C) and 26 ppm ($^{15}$N),
HN(CA)CO was carried out with 2048 ($^{1}$H) x 922 ($^{13}$C) x 72 ($^{15}$N) complex points and sweep widths of 12.98 ppm ($^{1}$H), 8 ppm ($^{13}$C) and 26 ppm ($^{15}$N),
B-HNCACB 2048 ($^{1}$H) x 128 ($^{13}$C) x 72 ($^{15}$N) complex points and sweep widths of 12.98 ppm ($^{1}$H), 60 ppm ($^{13}$C) and 26 ppm ($^{15}$N),
B-(H)N(CA)NH with 2048 ($^{1}$H) x 72 ($^{15}$N) x 72 ($^{15}$N) complex points and sweep widths of 12.98 ppm ($^{1}$H), and 26 ppm ($^{15}$N).
Nanog_aa1-85: interscan delay: 0.5 s
B-HNCO was carried out with 1024 ($^{1}$H) x 96 ($^{13}$C) x 92 ($^{15}$N) complex points and sweep widths of 12.98 ppm ($^{1}$H), 8 ppm ($^{13}$C) and 26 ppm ($^{15}$N),
B-HN(CA)CO: 1024 ($^{1}$H) x 88 ($^{13}$C) x 72 ($^{15}$N) complex points and sweep widths of 12.98 ppm ($^{1}$H), 8 ppm ($^{13}$C) and 26 ppm ($^{15}$N),
B-HNCACB: 1024 ($^{1}$H) x 96 ($^{13}$C) x 72 ($^{15}$N) complex points and sweep widths of 12.98 ppm ($^{1}$H), 60 ppm ($^{13}$C) and 22 ppm ($^{15}$N),

B-(H)N(CA)NH: 1024 ($^1$H) x 72 ($^{15}$N) x 72 ($^{15}$N) complex points and sweep widths of 12.98 ppm ($^1$H), and 26 ppm ($^{15}$N).
Oct4_aa1-145: interscan delay: between 0.12 and 0.2s
B-HNCO was carried out with 1536 ($^1$H) x 92 ($^{13}$C) x 72 ($^{15}$N) complex points and sweep widths of 16.6 ppm ($^1$H), 10 ppm ($^{13}$C) and 24 ppm ($^{15}$N),
B-HN(CA)CO: 1536 ($^1$H) x 92 ($^{13}$C) x 64 ($^{15}$N) complex points and sweep widths of 16.6 ppm ($^1$H), 10 ppm ($^{13}$C) and 24 ppm ($^{15}$N),
B-HNCACB: 1536 ($^1$H) x 128 ($^{13}$C) x 64 ($^{15}$N) complex points and sweep widths of 16.6 ppm ($^1$H), 65 ppm ($^{13}$C) and 24 ppm ($^{15}$N),
B-(H)N(CA)NH: 1536 ($^1$H) x 64 ($^{15}$N) x 64 ($^{15}$N) complex points and sweep widths of 16.6 ppm ($^1$H), and 24 ppm ($^{15}$N).
Sox2_aa115-236 (together with Sox2_aa1-187): interscan delay: between 0.12 and 0.2 s.
B-HNCO was carried out with 1536 ($^1$H) x 92 ($^{13}$C) x 64 ($^{15}$N) complex points and sweep widths of 16.6 ppm ($^1$H), 10 ppm ($^{13}$C) and 25 ppm ($^{15}$N),
B-HN(CA)CO: 1536 ($^1$H) x 64 ($^{13}$C) x 64 ($^{15}$N) complex points and sweep widths of 16.6 ppm ($^1$H), 10 ppm ($^{13}$C) and 26 ppm ($^{15}$N),
B-HNCACB: 1536 ($^1$H) x 84 ($^{13}$C) x 64 ($^{15}$N) complex points and sweep widths of 16.6 ppm ($^1$H), 65 ppm ($^{13}$C) and 26 ppm ($^{15}$N),
B-(H)N(CA)NH: 1536 ($^1$H) x 32 ($^{15}$N) x 32 ($^{15}$N) complex points and sweep widths of 16.6 ppm ($^1$H), and 24 ppm ($^{15}$N).
His6-AviTag-Sox2_aa234-317: interscan delay: between 0.12 and 0.2 s.
B-HNCO was carried out with 1536 ($^1$H) x 92 ($^{13}$C) x 72 ($^{15}$N) complex points and sweep widths of 16.6 ppm ($^1$H), 11 ppm ($^{13}$C) and 24 ppm ($^{15}$N),
B-HN(CA)CO: 1536 ($^1$H) x 92 ($^{13}$C) x 64 ($^{15}$N) complex points and sweep widths of 16.6 ppm ($^1$H), 11 ppm ($^{13}$C) and 24 ppm ($^{15}$N),
B-HNCACB: 1536 ($^1$H) x 128 ($^{13}$C) x 64 ($^{15}$N) complex points and sweep widths of 16.6 ppm ($^1$H), 65 ppm ($^{13}$C) and 24 ppm ($^{15}$N),
B-(H)N(CA)NH: 1536 ($^1$H) x 64($^{15}$N) x 64 ($^{15}$N) complex points and sweep widths of 16.6 ppm ($^1$H), and 24 ppm ($^{15}$N).
Sox2_aa115-317_C265A: d1=0.2 s.
B-HNCO: 2126 ($^1$H) x 128 ($^{13}$C) x 128 ($^{15}$N) complex points and sweep widths of 14 ppm ($^1$H), 7 ppm ($^{13}$C) and 22 ppm ($^{15}$N).
B-HN(CA)CO: 2126 ($^1$H) x 92 ($^{13}$C) x 92 ($^{15}$N) complex points and sweep widths of 14 ppm ($^1$H), 7 ppm ($^{13}$C) and 22 ppm ($^{15}$N). Non-uniform sampling at 35 %.
B-HNCACB : 2126 ($^1$H) x 128 ($^{13}$C) x 128 ($^{15}$N) complex points and sweep widths of 14 ppm ($^1$H), 60 ppm ($^{13}$C) and 22 ppm ($^{15}$N). Non-uniform sampling at 35 %.
B-(H)N(CA)NH : 18218($^1$H) x 92 ($^{15}$N) x 92 ($^{15}$N) complex points and sweep widths of 14 ppm ($^1$H), and 22 ppm ($^{15}$N). Non-uniform sampling at 35 %.

Spectra were processed with linear prediction of 16 or 32 complex points in both $^{13}$C, and $^{15}$N dimensions, cosine apodization in $^1$H and $^{15}$N dimensions, no apodization in $^{13}$C dimension, and zero filling to 2048, 512 and 256 complex points in $^1$H, $^{13}$C, and $^{15}$N dimensions, respectively. Assignment 2D $^1$H-$^{15}$N HSQC spectra were recorded using at least 1536 ($^1$H) x 256 ($^{15}$N) complex points and sweep widths of 16.23 ppm ($^1$H) and 30 ppm ($^{15}$N), and processed with zero filling to 4K and 1K in the proton and nitrogen dimensions, respectively.
$^1$H$_N$/$^{15}$N/$^{13}$C$\alpha$/$^{13}$C$\beta$/$^{13}$CO NMR assignments of OSNE peptides, together with the corresponding experimental details, have been deposited in the Biological Magnetic Resonance Data Bank (BMRB), with the accession numbers: 51534 (Sox2_aa1-42), 51717 (Esrrb_aa1-102), 51756 (Oct4_aa1-145), 51758 (Oct4_aa286-360), 51782 (His6-AviTag-Sox2_aa234-317_C265A), 51780 (Nanog_aa185).

#### 1.4. NMR monitoring of phosphorylation reactions and production of phosphorylated peptides

Phosphorylation reactions were carried out using $^{15}$N-labeled IDRs at 50 µM, in Hepes 20 mM, NaCl 50 mM, DTT or TCEP at 4 mM, ATP 1.5 mM, $MgCl_2$ at 5 mM, protease inhibitors (Roche), 7.5% $D_2O$, pH6.8 at 25°C in 100 µL using 3 mm diameter Shigemi tubes. We monitored the phosphorylation kinetics by recording time series of $^1$H-$^{15}$N SOFAST-HMQC spectra on a 600 MHz Bruker Avance II or a 700 MHz Bruker Avance Neo spectrometer, both equipped with cryogenically cooled triple resonance $^1$H[$^{13}$C/$^{15}$N] probes optimized for $^1$H detection.

The kinase was spiked in the IDR sample on ice just before filling the NMR tube, which was immediately placed in the spectrometer. About 2 minutes were necessary to reach a temperature equilibrium. The automatic shimming procedure was then executed, and short 1D $^{1}H$ and $^{1}H(^{15}N$-filtered)-SOFAST-HMQC spectra were recorded before the 2D spectra.
We recorded 2D spectra during the phosphorylation reactions as follows: 2D $^{1}H$-$^{15}N$ SOFAST-HMQC experiments were recorded using 2048 ($^{1}H$) x 96 ($^{15}N$) complex points and sweep widths of 16.6 ppm ($^{1}H$) and 26 ppm ($^{15}N$), 128 scans and interscan delays of 0.04 s; hence, the acquisition of one spectrum took 30 minutes. All spectra were processed zero filling to 2K and 1K in the direct and indirect dimensions, respectively. No apodization was applied for $^{1}H$-$^{15}N$ SOFAST-HMQC spectra.
After processing spectra in Topspin3, we measured peak intensities in NMRFAM-SPARKY [1]. Peaks were centered in every spectrum to follow peak shifting because of pH drifts. Progress curves were plotted and fitted in Kaleidagraph 4.5. In the case of the phosphorylation reactions that were not complete, we used decay curves to normalize phosphorylation build-up curves. At the opposite, for the phosphorylation reactions reaching ~100%, we used the phospho-peaks intensities for normalizing the build-up curves. Detailed descriptions of the methods can be found in previous reports and published protocols [2–5].

##### 1.5. Pull-down assays for interactomic analysis

Mouse Embryonic Stem Cells (mESCs) were harvested using a classical trypsin treatment (2x 150 $cm^2$ cell culture dishes, 70% confluency). The production of nuclear extracts was inspired by the procedures previously published by Gingras and colleagues [6]. Trypsin was blocked, and cell pellets were washed three times in PBS, before being resuspended in a first gentle lysis buffer containing HEPES at 10 mM, KCl at 10mM, EDTA at 0.5 mM, DTT at 1 mM, PMSF at 0.5 mM, 1% v/v NP40. After 10 minutes on ice, the cells were centrifuged 10 minutes at 15,000g. The supernatant containing the cytosolic fraction was discarded, and the pellets containing the nuclei were resuspended in a second lysis buffer containing HEPES at 20 mM, KCl at 250 mM, EDTA at 0.5 mM, DTT at 1 mM, PMSF at 0.5 mM, phosphatase inhibitors (2x PhosSTOP, Roche), 5% v/v glycerol, supplemented with 2 µL of benzonase (>250 units/µL, E1014 Sigma-Aldrich), before being sonicated on ice using a microtip sonicator and 3 pulses of 10 seconds. The lysis of nuclei was verified visually under the microscope using a cell counting chamber. The extract concentration used for the pull-downs was about 5 mg/mL, as measured by Bradford protein assay.
The pull-down assays were executed using 25 µL of streptavidin-coated magnetic beads (Magbeads streptavidine, Genscript), i.e. 50 µL of resuspended beads in the storing buffer. After every step described below, the tubes were placed on a magnetic rack to collect the beads, while the supernatant was removed with a pipette. The fresh beads were washed 3 times during 5 minutes in 500 µL of a PBS buffer. 1 nmol of biotinylated (using BirA, see above) AviTag-chimera peptides (either AviTag-Sox2(aa115-240), AviTag-Sox2(aa234-317_C265A), phospho-AviTag-Sox2(aa115-240), or phospho-AviTag-Sox2(aa234-317_C265A)) were diluted in 500 µL of PBS and incubated with the beads during one hour at room temperature under rotary agitation. The supernatant was then removed, the beads were washed 3 times during 5 minutes in 500 µL of a PBS buffer.
The mESCs extract (200 µL, generated from 15 million cells) was mixed with the beads, and then incubated during one hour at room temperature under rotary agitation. The supernatant was removed and the beads were washed 3 times during 5 minutes in 500 µL of a buffer containing HEPES at 20 mM, KCl at 250 mM, EDTA at 0.5 mM, DTT at 1 mM, PMSF at 0.5 mM + PhosSTOP 1x.

##### 1.6. Mass spectrometry-based proteomics analysis of pull-down assays

*Sample Preparation:* The beads were resuspended in 100 µL of 25 mM $NH_4HCO_3$ and digested by adding 0.2 µg of trypsin/LysC (Promega) for 1 h at 37 °C. Samples were then loaded into custom-made C18 StageTips packed by stacking three AttractSPE® disk (#SPE-Disks-Bio-C18-100.47.20 Affinisep) into a 200 µL micropipette tip for desalting. Peptides were eluted using a ratio of 40:60 $CH_3CN$:$H_2O$ + 0.1% formic acid and vacuum concentrated to dryness with a SpeedVac apparatus. Peptides were reconstituted in 10 µL of injection buffer in 0.3% trifluoroacetic acid (TFA) before liquid chromatography-tandem mass spectrometry (LC-MS/MS) analysis.
*LC-MS/MS Analysis:* Online chromatography was performed with an RSLCnano system (Ultimate 3000, Thermo Scientific) coupled to an Orbitrap Fusion Tribrid mass spectrometer (Thermo Scientific). Peptides were trapped on a C18 column (75 µm inner diameter × 2 cm; nanoViper Acclaim PepMap™ 100, Thermo Scientific) with buffer A (2/98 MeCN/$H_2O$ in 0.1% formic acid) at a flow rate of 4.0 µL/min over 4 min. Separation was performed on a 50 cm x 75 µm C18 column (nanoViper Acclaim PepMap™ RSLC, 2 µm, 100 Å, Thermo Scientific)

regulated to a temperature of 55 °C with a linear gradient of 5% to 25% buffer B (100% MeCN in 0.1% formic acid) at a flow rate of 300 nL/min over 100 min. Peptides were ionized by a nanospray ionization (NSI) ion source at 2.2 kV. Full-scan MS in the Orbitrap was set at a scan range of 400-1500 with a resolution at 120,000 and ions from each full scan were fragmented in higher-energy collisional dissociation mode (HCD) and analyzed in the linear ion trap in rapid mode. The fragmentation was set top speed mode in data-dependent analysis (DDA). We selected ions with charge state from 2+ to 7+ for screening. Normalized collision energy (NCE) was set to 30, AGC target to 20,000 and the dynamic exclusion to 30s.
*Data analysis:* For identification, the datasets were searched against the *Mus Musculus* (UP000000589) UniProt database using Sequest HT through proteome discoverer (version 2.2). Enzyme specificity was set to trypsin and a maximum of two miss cleavages sites were allowed. Oxidized methionine, phosphorylation of serines, threonines and tyrosines, carbamidomethylation of cysteines and N-terminal acetylation were set as variable modifications. Maximum allowed mass deviation was set to 10 ppm for monoisotopic precursor ions and 0.6 Da for MS/MS peaks. The resulting files were further processed using myProMS v3.9.3 (https://github.com/bioinfo-pf-curie/myproms; Poullet et al., 2007 [7]). False-discovery rate (FDR) was calculated using Percolator [8] and was set to 1% at the peptide level for the whole study. Label-free quantification was performed using peptide extracted ion chromatograms (XICs), computed with MassChroQ [9] v.2.2.1. For protein quantification, XICs from proteotypic peptides shared between compared conditions (TopN matching) were used, missed cleavages and peptide modifications were not allowed. Median and scale normalization at peptide level was applied on the total signal to correct the XICs for each biological replicate (N=2). To estimate the significance of the change in protein abundance, a linear model (adjusted on peptides and biological replicates) was performed, and p-values were adjusted using the Benjamini–Hochberg FDR procedure.
The mass spectrometry proteomics raw data have been deposited to the ProteomeXchange Consortium via the PRIDE partner repository dataset [10]: identifier PXD 040573 ( & **Password:** sVM686zJ).

#### 1.7. Recombinant production of Pin1 and NMR analysis of its interaction with Sox2 or phospho-Sox2

The plasmid containing the gene coding for the Pin1-WW domain was a kind gift from Isabelle Landrieu. The production was executed according to the previously published protocol [11]. The NMR analysis of binding with phosphoSox2(aa115-240) were performed with the GST-Pin1-WW construct and $^{15}$N-labeled Sox2(aa115-240) mixed in stoichiometric proportions, either at 50 or 10 μM for non-phospho and phosphoSox2, respectively. The solution contained Hepes at 20 mM, NaCl at 50 mM, urea at 0.25 mM (left-overs from Sox2(aa115-240) stock, stored at 2 M urea for solubility, see above), 5% $D_2O$ and DSS at 0.1 mM, at pH=7.0. The 2D $^{1}$H-$^{15}$N SOFAST-HMQC spectra were recorded at 283K, using a 600 MHz Bruker Avance II equipped with a cryogenically cooled triple resonance $^{1}$H[$^{13}$C/$^{15}$N] probe and a 5 mm diameter Shigemi tube,
The experiments were recorded using 1536 ($^{1}$H) x 128 ($^{15}$N) complex points and sweep widths of 16.0 ppm ($^{1}$H) and 264ppm ($^{15}$N), 64 or 128 scans and interscan delays of 0.04 s. The spectra were processed with zero filling to 2K and 1K in the direct and indirect dimensions, respectively. Cosine apodization was applied in both dimensions. After processing spectra in Topspin3, we measured peak intensities in NMRFAM-SPARKY [1], and plotted the intensity ratios in Kaleidagraph 4.5.

#### 2. Supplementary Figures

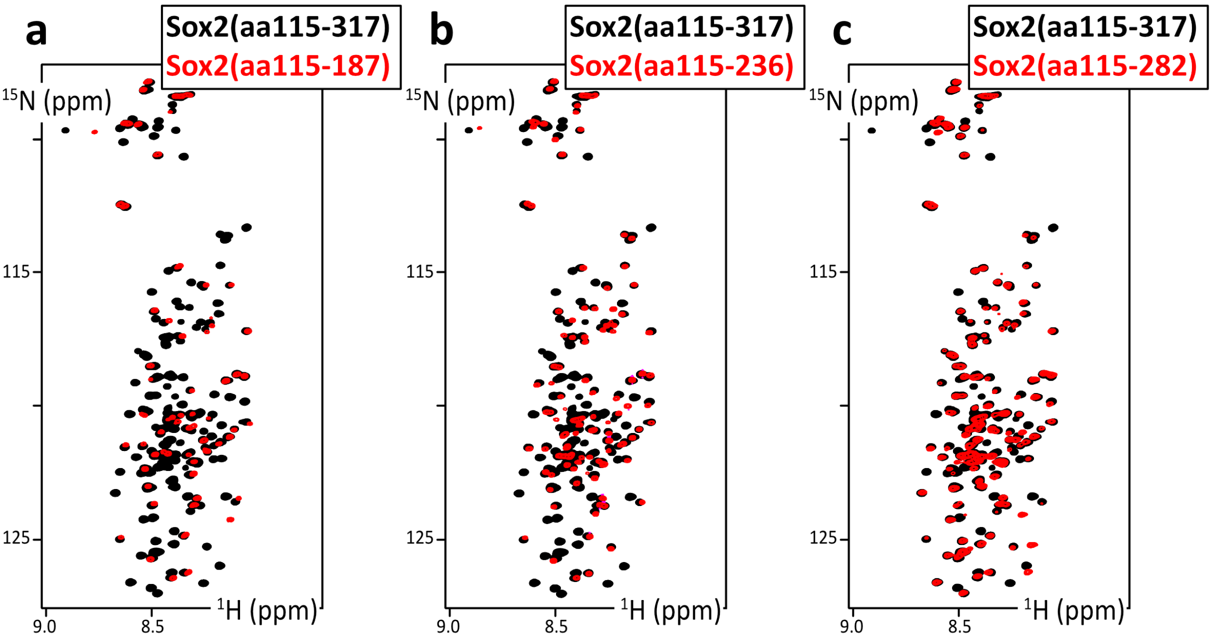


**Figure S1:** *Overlays of 2D $^{1}H$-$^{15}N$ HSQC spectra of Sox2(aa115-317_C265A) and Sox2(aa115-187), Sox2(aa115-236) and Sox2(aa115-282_C265A). These spectra have been recorded in a buffer containing urea at 0.25 M, except for Sox2(aa115-187), at 283K and 700 MHz.*

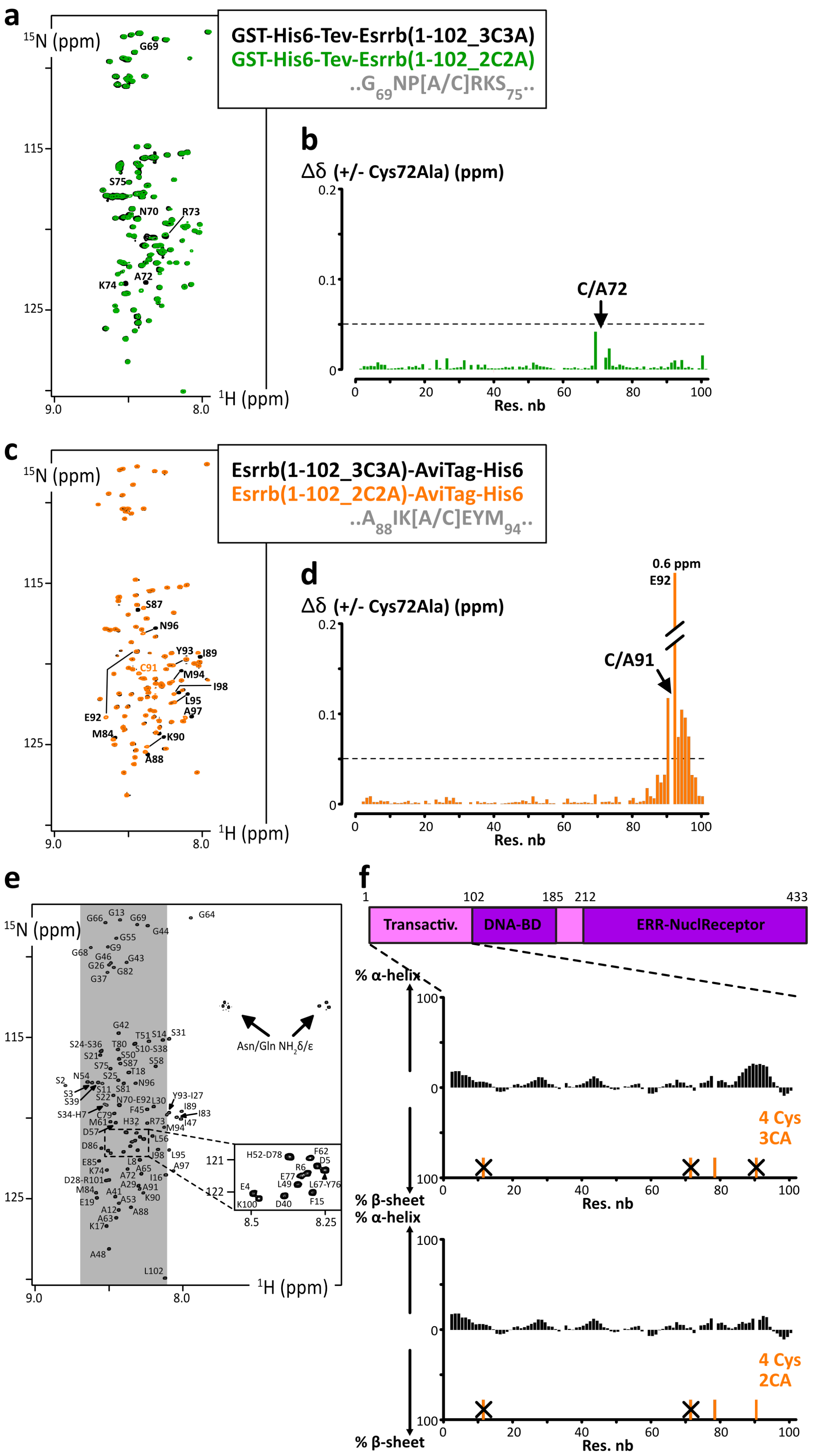

a
$^{15}$N (ppm)
GST-His6-Tev-Esrrb(1-102_3C3A)
GST-His6-Tev-Esrrb(1-102_2C2A)
..G69NP[A/C]RKS75..
G69
S75
N70
R73
A72
K74
115
125
9.0
8.0
$^{1}$H (ppm)
b
Δδ (+/- Cys72Ala) (ppm)
0.2
0.1
0
C/A72
0
20
40
60
80
100
Res. nb
c
$^{15}$N (ppm)
Esrrb(1-102_3C3A)-AviTag-His6
Esrrb(1-102_2C2A)-AviTag-His6
..A88IK[A/C]EYM94..
S87
N96
Y93
I89
C91
M94
I98
L95
A97
E92
M84
K90
A88
115
125
9.0
8.0
$^{1}$H (ppm)
d
0.6 ppm
E92
Δδ (+/- Cys72Ala) (ppm)
0.2
C/A91
0.1
0
0
20
40
60
80
100
Res. nb
e
$^{15}$N (ppm)
115
125
9.0
8.0
$^{1}$H (ppm)
Asn/Gln NH$_2$δ/ε
G13
G66
G69
G64
G44
G55
G9
G68
G46
G43
G26
G82
G37
G42
T51
S14
S31
S24-S36
T80
S10-S38
S21
S50
S75
S87
S58
T18
S2
N54
S25
N96
S3
S11
S81
S39
S22
N70-E92
L30
Y93-I27
I89
S34-H7
C79
F45
I83
M61
H32
R73
I47
D57
M94
L56
D86
I98
L95
E85
L8
A97
K74
A65
D28-R101
A72
I16
M84
A41
A29
A91
E19
A53
K90
A12
A88
A63
K17
A48
L102
121
122
H52-D78
F62
R6
D5
E77
L49
L67-Y76
E4
K100
D40
F15
8.5
8.25
f
1
102
185
212
433
Transactiv.
DNA-BD
ERR-NuclReceptor
% α-helix
100
0
100
% β-sheet
4 Cys
3CA
0
20
40
60
80
100
Res. nb
% α-helix
100
0
100
% β-sheet
4 Cys
2CA
0
20
40
60
80
100
Res. nb

**Figure S2: a.** *Overlay of 2D $^1H$-$^{15}N$ HSQC spectra of Esrrb(aa1-102_C12A-C72A-C91A) (black) and Esrrb(aa1-102_C12A-C91A) (green);* ***b.*** *Chemical shift perturbations between the two constructs in* ***a.*** *using $\Delta\delta=[(\Delta\delta_{1H}{}^2+(\Delta\delta_{15N}/5)^2)/2]^{1/2}$;* ***c.*** *Overlay of 2D $^1H$-$^{15}N$ HSQC spectra of Esrrb(aa1-102_C12A-C72A-C91A) (black) and Esrrb(aa1-102_C12A-C72A) (orange);* ***d.*** *Chemical shift perturbations between the two constructs in* ***a.***; ***e.*** *2D $^1H$-$^{15}N$ HSQC spectrum of the N-terminal IDRs of human Esrrb(aa1-102_C12A-C72A-C91A), the labels indicating the assignments;* ***f.*** *Primary structure of human Esrrb, and Secondary structure propensities of Esrrb(aa1-102_C12A-C72A-C91A) and Esrrb(aa1-102_C12A-C72A) calculated from the experimental chemical shifts of the peptide backbone $C\alpha$ and $C\beta$, using the ncSPC algorithm [12,13].*

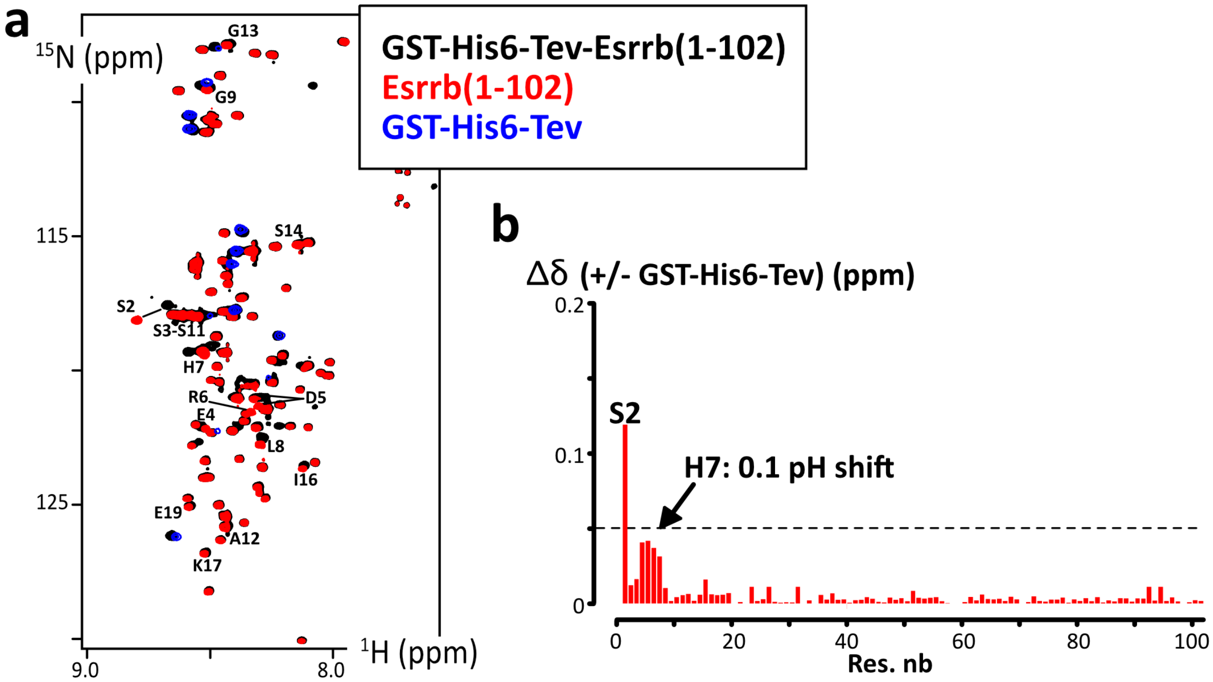


**Figure S3:** ***a.*** *Overlay of 2D $^{1}H$-$^{15}N$ HSQC spectra of GST-His6-Tev-Esrrb(aa1-102_C12A-C72A-C91A) (black), Esrrb(aa1-102_C12A-C72A-C91A) (red) and GST-His6-Tev (blue);* ***b.*** *Chemical shift perturbations between the GST-His6-Tev-Esrrb(aa1-102_C12A-C72A-C91A) and Esrrb(aa1-102_C12A-C72A-C91A) after TEV cleavage, using $\Delta\delta=[(\Delta\delta_{1H}{}^{2}+(\Delta\delta_{15N}/5)^{2})/2]^{1/2}$.*

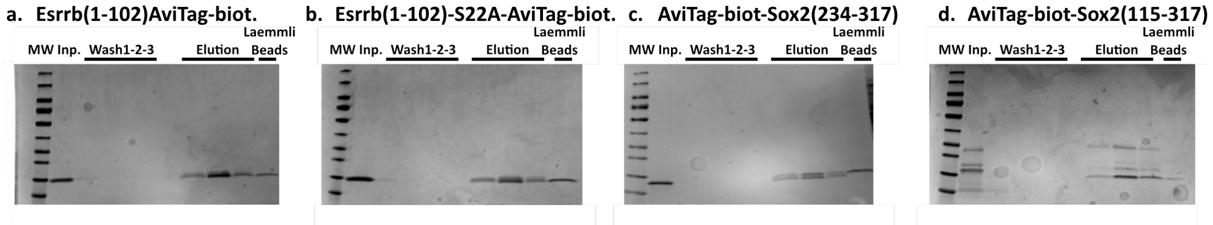


***Figure S4 :*** *SDS-PAGE analysis of binding assays : 1 nmol of biotinylated AviTag-IDRs (Input : Inp.) were incubated with 25 mL of streptavidin-coated magnetic beads ; the 3 washing steps showed the stability of the binding ; the elutions were performed with a Laemmli buffer, which provokes also the release of the streptavidine, whose band is shown in the last lane and unfortunately overlaps with the AviTag-IDRs constructs here.* ***d.*** *We show here one of our tests with a batch of AviTag-Sox2(aa115-317_C265A), which was partially proteolyzed; this is to show that one of the peptides that do not contain the AviTag is removed by the first wash.*

### 3. Supplementary material 3: Protein sequences

## 3.1. GB1

QYKLILNGKTLKGETTTEAVDAATAEKVFKQYANDNGVDGEWTYDDATKTFTVTEGG

#### 3.2. Oct4

##### 3.2.1. Sequence alignment: mammals

```
S: Serine
P : Proline
SP : phosphorylation motif for MAPKand Cdk
FILVYW : hydrophobic
DE : Asp/Glu
KR : Lys/Arg
T : Thr
XXX : DND-BD in crystal (3L1P)

oct4-human        MAGHLASDFAFSPPPGGGGDGPGGPEPGWVDPRTWLSFQGPPGGPGIGPGVGPGSEVWGI 60
oct4-mice         MAGHLASDFAFSPPPGGG-DGSAGLEPGWVDPRTWLSFQGPPGG----PGIGPGSEVLGI 55
oct4-cow          MAGHLASDFAFSPPPGGGGDGPGGPEPGWVDPRTWMSFQGPPGGSGIGPGVVPGAEVWGL 60
oct4-dog          MAGHLASDLAFSPSPGGGGDGPGGPDPGWGDPRAWLSFPGPPGGPALGPGVGPGAEVWGL 60
Oct4-sheep        MAGHLASDFAFSPPPGGGGDGPGGPEPGWVDPRTWMSFQGPPGGSGIGPGVVPGAEVWGL 60
pou5f1-spermwhale MAGHLASDFAFSPPPGGGGDGPGGPEPGWVDPRTWMSFQGPPGGSGIGPGVGPGAEVWGL 60
                  ********:****.**** *...* :*** ***:*:** ***.*    **: **:** *:

oct4-human        PPCPPPYEFCGGMAYCGPQVGVGLVPQGGLETSQPEGEAGVGVESNSDGASPEPCTVTPG 120
oct4-mice         SPCPPAYEFCGGMAYCGPQVGLGLVPQVGVETLQPEGQAGARVESNSEGTSSEPCADRPN 115
oct4-cow          PPCPPPYDLCGGMAYCAPQVGVGPVPPGGLETPQPEGEAGAGVESNSEGASPDPCAAPAG 120
oct4-dog          PPCPPPYEFCGGMAYCGPQVGVGLLPQGGLDTSQPEGERGAGLEGSSEGASPEPCAAPPG 120
Oct4- sheep       PPCPPPYDLCGGMAYCAPQVGVGPVPPGGLETPQPEGEAGAGVESNSEGASPDPCAAPAG 120
pou5f1-spermwhale PACPPPYDLCGGMAYCAPQVGVGLVPQGGLETPQPEGEAGAGVESKSEGASPEPCAAPAG 120
                  ..***.*::****:**.****:* :*  *::* ****: *. :*..*:*:*. **:  ..

oct4-human        AVKLEKEKLEQNPEESQDIKALQKELEQFAKLLKQKRITLGYTQADVGLTLGVLFGKVFS 180
oct4-mice         AVKL--EKVEPTPEESQDMKALQKELEQFAKLLKQKRITLGYTQADVGLTLGVLFGKVFS 173
oct4-cow          APKLDKEKLEPNPEESQDIKALQKDLEQFAKLLKQKRITLGYTQADVGLTLGVLFGKVFS 180
oct4-dog          VVKPDKEKLEQNPEESQDIKALQKDLEQFAKLLKQKRITLGYTQADVGLTLGVLFGKVFS 180
Oct4-sheep        AAKLDKEKLEPNPEESQDIKALQKDLEQFAKLLKQKRITLGYTQADVGLTLGVLFGKVFS 180
pou5f1-spermwhale AEKVDKEKLEPNPEESQDIKALQKDLEQFAKLLKQKRITLGYTQADVGLTLGVLFGKVFS 180
                  . *   **:* ..*****:*****:***********************************

oct4-human        QTTICRFEALQLSFKNMCKLRPLLQKWVEEADNNENLQEICKAETLVQARKRKRTSIENR 240
oct4-mice         QTTICRFEALQLSLKNMCKLRPLLEKWVEEADNNENLQEICKSETLVQARKRKRTSIENR 233
oct4-cow          QTTICRFEALQLSFKNMCKLRPLLQKWVEEADNNENLQEICKAETLVQARKRKRTSIENR 240
oct4-dog          QTTICRFEALQLSFKNMCKLRPLLQKWVEEADNNENLQEICKAETLVQARKRKRTSIENR 240
Oct4-sheep        QTTICRFEALQLSFKNMCKLRPLLQKWVEEADNNENLQEICKAETLVQARKRKRTSIENR 240
pou5f1-spermwhale QTTICRFEALQLSFKNMCKLRPLLQKWVEEADNNENLQEICKAETLMQARKRKRTSIENR 240
                  *************:**********:*****************:***:*************

oct4-human        VRGNLENLFLQCPKPTLQQISHIAQQLGLEKDVVRVWFCNRRQKGKRSSSDYAQREDFEA 300
oct4-mice         VRWSLETMFLKCPKPSLQQITHIANQLGLEKDVVRVWFCNRRQKGKRSSIEYSQREEYEA 293
oct4-cow          VRGNLESMFLQCPKPTLQQISHIAQQLGLEKDVVRVWFCNRRQKGKRSSSDYSQREDFEA 300
oct4-dog          VRGNLENMFLQCPKPTLQQISHIAQQLGLEKDVVRVWFCNRRQKGKRSSSDYSQREDFEA 300
Oct4-sheep        VRGNLESMFLQCPKPTLQQISHIAQQLGLEKDVVRVWFCNRRQKGKRSSSDYSQREDFEA 300
pou5f1-spermwhale VRGNLESMFLQCPKPTLQQISHIAQQLGLEKDVVRVWFCNRRQKGKRSSSDYSQREDFEA 300
                  ** .**.:**:****:****: **:*****:*****************  :*:***::**

oct4-human        AGSPFSGGPVSFPLAPGPHFGTPGYGSPHFTALYSSVPFPEGEAFPPVSVTTLGSPMHSN 360
oct4-mice         TGTPFPGGAVSFPLPPGPHFGTPGYGSPHFTTLYS-VPFPEGEAFPSVPVTALGSPMHSN 352
oct4-cow          AGSPFTGGPVSSPLAPGPHFGTPGYGGPHFTTLYSSVPFPEGEVFPSVSVTALGSPMHAN 360
oct4-dog          AGSPFSGAPVSFPLAPGPHFGTPGYGGPHFTTLYSSVPLPEGEGFPSVSVTTLGSPMHSN 360
Oct4-sheep        AGSPFAGGPVSFPLAPGPPFGTPGYGGPHFTTLYSSVPFPEGEAFPSVSVTALGSPMHAN 360
pou5f1-spermwhale AGSPFSGGPVSFPLAPGPHFGTPGYGGPHFTTLYSSVPFPEGEAFPSVSVTTLGSPMHSN 360
                  :*.**.*..** **.*** **:****.****:*** **:**** **.*.**.******:*
```

>oct4-human
MAGHLASDFAFSPPPGGGGDGPGGPEPGWVDPRTWLSFQGPPGGPGIGPGVGPGSEVWGIPPCPPPYEFC
GGMAYCGPQVGVGLVPQGGLETSQPEGEAGVGVESNSDGASPEPCTVTPGAVKLEKEKLEQNPEESQDIK
ALQKELEQFAKLLKQKRITLGYTQADVGLTLGVLFGKVFSQTTICRFEALQLSFKNMCKLRPLLQKWVEE
ADNNENLQEICKAETLVQARKRKRTSIENRVRGNLENLFLQCPKPTLQQISHIAQQLGLEKDVVRVWFCN
RRQKGKRSSSDYAQREDFEAAGSPFSGGPVSFPLAPGPHFGTPGYGSPHFTALYSSVPFPEGEAFPPVSV
TTLGSPMHSN

##### 3.2.2. Sequence alignment: vertebrates

```
oct4-human          MAGHLASDFAFS----------PPPGGGGDGPGGPEPGWVDP------------------ 32
pou5f1-Danio        MTERAQSPTAADCRPYEVNRAMYPQAAGLDGLGGASLQFAHGMLQDPSLIFNKAHFNGIT 60
oct4-Gallus         MHVKAKN--------------LLRMCKWLKGLRNA------------------------- 21
                    *  :  .                      .*  ..

oct4-human          ----RTWLSFQGPPGGPGIGPG-------------VGPGSEVWGIPPCPPPYEFCGG--- 72
pou5f1-Danio        PATAQTFFPFSGDFKTNDLQGGDFTQPKHWYPFAAPEFTGQVAGATAATQPANISPPIGE 120
oct4-Gallus         --------------------------------------RGSTWGRSGGRKPMRSSG---- 39
                                                           ... * .    * . .

oct4-human          -----MAYCGPQVGVGLVPQGGLETSQP--------EGEAGVGVESNSDG---------- 109
pou5f1-Danio        TREQIKMPSEVKTEKDVEEYGNEENKPPSQYHLTAGTSSVPTGVNYYTPWNPNFWPGLSQ 180
oct4-Gallus         -----RLPRSADPG-----WGNHANR----------AAVVTRGISSHSPR---------- 69
                               .        *.  .            . .  *:.  :

oct4-human          ----------------------------------------ASPEPCTVTPGAVKLEKEKL 129
pou5f1-Danio        ITAQANISQAPPTPSASSPSLSPSPPGNGFGSPGFFSGGTAQNIPSAQAQSAPRSSGSSS 240
oct4-Gallus         -------------------------------------------VCLCLCQDAP------- 79
                                                                      .*

oct4-human          EQNPEESQDIKALQKELEQFAKLLKQKRITLGYTQADVGLTLGVLFGKVFSQTTICRFEA 189
pou5f1-Danio        GGCSDSEEEETLTTEDLEQFAKELKHKRITLGFTQADVGLALGNLYGKMFSQTTICRFEA 300
oct4-Gallus         ------------TSEELEQFAKDLKHKRIMLGFTQADVGLALGTLYGKMFSQTTICRFEA 127
                                  ::****** **:*** **:*******:** *:**:***********

oct4-human          LQLSFKNMCKLRPLLQKWVEEADNNENLQEICKAE-TLVQARKRKR-TSIENRVRGNLEN 247
pou5f1-Danio        LQLSFKNMCKLKPLLQRWLNEAENSENPQDMYKIERVFVDTRKRKRRTSLEGTVRSALES 360
oct4-Gallus         LQLSFKNMCKLKPLLQRWLNEAENTDNMQEMCNAEQVLAQARKRKRRTSIETNVKGTLES 187
                    ***********:****:*::**:*.:* *:: : * .:.::***** **:*  *:. **.

oct4-human          LFLQCPKPTLQQISHIAQQLGLEKDVVRVWFCNRRQKGKRSSSDYAQREDFEAAGSPFSG 307
pou5f1-Danio        YFVKCPKPNTLEITHISDDLGLERDVVRVWFCNRRQKGKRLALPFDDECVEAQYYEQSPP 420
oct4-Gallus         FFRKCVKPSPQEISQIAEDLNLDKDVVRVWFCNRRQKGKRLLLPFGNESEGVMYDMNQSL 247
                     * :* **.  :*::*:::*.*::****************    : :.          .

oct4-human          GPVSFPLAPGPHFGTPGYGSPHFTALYSSVPFPEGEAFPPVSVTTLGSPMHSN 360
pou5f1-Danio        PPPHMGGTVLPGQGYPGPAHPGGAPALYMPSLHRPDVFKNGLHPGLVGHLTS- 472
oct4-Gallus         VPPGLP-IPVTSQGYS----LAPSPPVYMPPFHKAEMFPPPLQPGISMNNSSH 295
                     *  :     .  * .       :.     .: . : *     . :     *
```

##### 3.2.3. Disorder prediction

https://st-protein.chem.au.dk/odinpred
https://www.nature.com/articles/s41598-020-71716-1

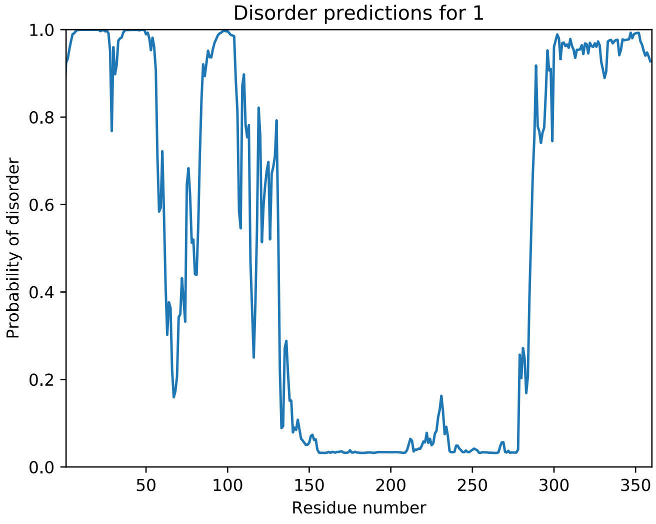


##### 3.2.4. Coding DNA sequences, produced protein constructs

**Oct4-aa1-145**

Synthesized sequence:

```
Ccgcgggtgagaacctgtacttccaaggcatggcgggtcacctggcgagcgattttgcgtttagcccgccgccgggtggtggtggtgacggtcc
gggtggcccggaaccgggttgggtggatccgcgtacctggctgagcttccaaggtccgccgggtggcccgggtattggtccgggtgtgggcccg
ggtagcgaggtttggggtattccgccgtgcccgccgccgtacgaattttgcggtggcatggcgtattgcggtccgcaagtgggcgttggtctgg
ttccgcaaggtggcctggaaaccagccagccggagggtgaagcgggcgtgggtgttgagagcaacagcgatggtgcgagcccggaaccgtgcac
cgtgaccccgggtgcggttaagctggagaaggaaaaactggagcagaacccggaggaaagccaagatatcaaggcgctgcagaaagaaaacctg
tactttcaaggtggcgcgggtggcgcgggtggccagtataaactgattctgaacggcaagaccctgaaaggtgaaaccaccaccgaagcggtgg
```

atgcggcgaccgctgagaaggttttcaaacagtacgcgaacgacaacggcgtggatggcgagtggacctatgacgatgcgaccaagacctttac
cgttaccgaaggtggctaaaagctt

<u>Translates into:</u>
AGENLYFQGMAGHLASDFAFSPPPGGGGDGPGGPEPGWVDPRTWLSFQGPPGGPGIGPGVGPGSEVWGIPPCPPPYEFCGGMAYCGPQVGVGLV
PQGGLETSQPEGEAGVGVESNSDGASPEPCTVTPGAVKLEKEKLEQNPEESQDIKALQKENLYFQGGAGGAGGQYKLILNGKTLKGETTTEAVD
AATAEKVFKQYANDNGVDGEWTYDDATKTFTVTEGG*

<u>After TEV-cleavage *(leaving 1 Gly in N-ter, and ENLYFQ in C-ter)*:</u>

```
          10         20         30         40         50         60
G MAGHLASDFA FSPPPGGGGD GPGGPEPGWV DPRTWLSFQG PPGGPGIGPG VGPGSEVWGI

        70         80         90        100        110        120
PPCPPPYEFC GGMAYCGPQV GVGLVPQGGL ETSQPEGEAG VGVESNSDGA SPEPCTVTPG

       130        140
AVKLEKEKLE QNPEESQDIK ALQKENLYFQ
```

### Oct4-aa286-360

<u>Synthesized sequence:</u>
Ccgcgggtgagaacctgtactttcagggcaagcgtagcagcagcgactatgcgcaacgtgaggatttcgaagcggcgggtagcccgtttagcgg
tggcccggtgagcttcccgctggcgccgggtccgcactttggtaccccgggttatggcagcccgcacttcaccgcgctgtatagcagcgttccg
ttcccggagggtgaagcgtttccgccggtgagcgttaccaccctgggcagcccgatgcacagcaacgaaaatctgtactttcagggtggcgcgg
gtggcgcgggtggccaatataagctgatcctgaacggcaagaccctgaaaggcgaaaccaccaccgaagcggtggatgcggcgaccgctgagaa
ggtttttaaacagtacgcgaacgacaacggtgtggatggcgagtggacctatgacgatgcgaccaaaaccttcaccgttaccgaaggtggctaa
aagctt

<u>Translates into:</u>
AGENLYFQGKRSSSDYAQREDFEAAGSPFSGGPVSFPLAPGPHFGTPGYGSPHFTALYSSVPFPEGEAFPPVSVTTLGSPMHSNENLYFQGGAG
GAGGQYKLILNGKTLKGETTTEAVDAATAEKVFKQYANDNGVDGEWTYDDATKTFTVTEGG*

<u>After TEV-cleavage *(leaving 1 Gly in N-ter, and ENLYFQ in C-ter)*:</u>

```
    290        300
G KRSSS DYAQREDFEA

       310        320        330        340        350        360
AGSPFSGGPV SFPLAPGPHF GTPGYGSPHF TALYSSVPFP EGEAFPPVSV TTLGSPMHSN ENLYFQG
```

#### 3.3. Sox2

##### 3.3.1. Sequence alignment: mammals

S: Serine
P : Proline
**SP : phosphorylation motif for MAPKand Cdk**
FILVYW : hydrophobic
DE : Asp/Glu
KR : Lys/Arg
T : Thr
XXX : DND-BD in crystal (1GT0)

```
sox2-human               MYNMMETELKPPGPQQTSGGGGG-------NSTAAAAGGNQKNSPDRVKRPMNAFMVWSR 53
sox2-mus                 MYNMMETELKPPGPQQASGGGGGGG-----NATAAATGGNQKNSPDRVKRPMNAFMVWSR 55
sox2-Bos                 MYNMMETELKPPGPQQTSGGGGGGGG----NSTAAAAGGNQKNSPDRVKRPMNAFMVWSR 56
sox2-Canis               MYNMMETELKPPGPQQTSGGGGGGGGGGGGNSTAAAAGGNQKNSPDRVKRPMNAFMVWSR 60
sox2-Capra               MYNMMETELEQPGLQHNSGGGGGGGG----NSTAAAAGGNQKNSPDRVKRPMNAFMVWSR 56
sox2-Balaenoptera        ---------------------------------AAAAGGNQKNSPDRGKRPMNAFMVWSR 27
                                                          ***:********** ************

sox2-human               GQRRKMAQENPKMHNSEISKRLGAEWKLLSETEKRPFIDEAKRLRALHMKEHPDYKYRPR 113
sox2-mus                 GQRRKMAQENPKMHNSEISKRLGAEWKLLSETEKRPFIDEAKRLRALHMKEHPDYKYRPR 115
sox2-Bos                 GQRRKMAQENPKMHNSEISKRLGAEWKLLSETEKRPFIDEAKRLRALHMKEHPDYKYRPR 116
sox2-Canis               GQRRKMAQENPKMHNSEISKRLGAEWKLLSETEKRPFIDEAKRLRALHMKEHPDYKYRPR 120
sox2-Capra               GQRRKMAQENPKMHNSEISKRLGAEWKLLSETEKRPFIDEAKRLRALHMKEHPDYKYRPR 116
sox2-Balaenoptera        GQRRKMAQENPKMHNSEISKRLGAEWKLLSETEKRPFIDEAKRLRALHMKEHPDYKYRPR 87
                         *******************:****************************************

sox2-human               RKTKTLMKKDKYTLPGGLLAPGGNSMASGVGVGAGLGAGVNQRMDSYAHMNGWSNGSYSM 173
sox2-mus                 RKTKTLMKKDKYTLPGGLLAPGGNSMASGVGVGAGLGAGVNQRMDSYAHMNGWSNGSYSM 175
sox2-Bos                 RKTKTLMKKDKYTLPGGLLAPGGNSMASGVGVGAGLGAGVNQRMDSYAHMNGWSNGSYSM 176
sox2-Canis               RKTKTLMKKDKYTLPGGLLAPGGNSMASGVGVGAGLGAGVNQRMDSYAHMNGWSNGSYSM 180
sox2-Capra               RKTKTLMKKDKYTLPGGLLAPGGNIMASGVGVGAGLGAGVNQRMDSYAHMNGWSNGSYSM 176
sox2-Balaenoptera        RKTKTLMKKDKYRRAGGLLAPGGNSMASGVGVGAGLGAGVNQRMDSYAHMNGWSNGSYSM 147
                         ************  .********* ****************************** ****

sox2-human               MQDQLGYPQHPGLNAHGAAQMQPMHRYDVSALQYNSMTSSQTYMNGSPTYSMSYSQQGTP 233
sox2-mus                 MQEQLGYPQHPGLNAHGAAQMQPMHRYDVSALQYNSMTSSQTYMNGSPTYSMSYSQQGTP 235
sox2-Bos                 MQDQLGYPQHPGLNAHGAAQMQPMHRYDVSALQYNSMTSSQTYMNGSPTYSMSYSQQGTP 236
sox2-Canis               MQDQLGYPQHPGLNAHGAAQMQPMHRYDVSALQYNSMTSSQTYMNGSPTYSMSYSQQGTP 240
sox2-Capra               MQDQLGYPQHPGLNAHGAAQMQPMHRYDVSALQYNSMTSSQTYMNGSPTYSMSYSQQGTP 236
sox2-Balaenoptera        MQDQLGYPQHPGLNAHGAAQMQPMHRYDVSALQYNSMTSSQTYMNGSPTYSMSYSQQGTP 207
                         **:*************.*******************************************

sox2-human               GMALGSMGSVVKSEASSSPPVVTSSSHSRAPCQAGDLRDMISMYLPGAEVPEPAAPSRLH 293
sox2-mus                 GMALGSMGSVVKSEASSSPPVVTSSSHSRAPCQAGDLRDMISMYLPGAEVPEPAAPSRLH 295
sox2-Bos                 GMALGSMGSVVKSEASSSPPVVTSSSHSRAPCQAGDLRDMISMYLPGAEVPEPAAPSRLH 296
sox2-Canis               GMALGSMGSVVKSEASSSPPVVTSSSHSRAPCQAGDLRDMISMYLPGAEVPEPAAPSRLH 300
sox2-Capra               GMALGSMGSVVKSEASSSPPVVTSSSHSRPPCQAGDLRDMISMYLPGDEVPEPAAPSRLH 296
sox2-Balaenoptera        GMALGSMGSVVKSEASSSPPVVTSSSHSRAPCQAGDLRDMISMYLPGAEVPEPAAPSRLH 267
                         **:**************************.***************** ************

sox2-human               MSQHYQSGPVPGTAINGTLPLSHM 317
sox2-mus                 MAQHYQSGPVPGTAINGTLPLSHM 319
sox2-Bos                 MSQHYQSGPVPGTAINGTLPLSHM 320
sox2-Canis               MSQHYQSGPVPGTAINGTLPLSHM 324
sox2-Capra               MSQHYQSGAVPGTAINGILPLSHM 320
sox2-Balaenoptera        MSQHYQSGPVPGTAINGTLPLSHM 291
                         *:******.******** ******
```

```
>sox2-human
MYNMMETELKPPGPQQTSGGGGGNSTAAAAGGNQKNSPDRVKRPMNAFMVWSRGQRRKMAQENPKMHNSE
ISKRLGAEWKLLSETEKRPFIDEAKRLRALHMKEHPDYKYRPRRKTKTLMKKDKYTLPGGLLAPGGNSMA
SGVGVGAGLGAGVNQRMDSYAHMNGWSNGSYSMMQDQLGYPQHPGLNAHGAAQMQPMHRYDVSALQYNSM
TSSQTYMNGSPTYSMSYSQQGTPGMALGSMGSVVKSEASSSPPVVTSSSHSRAPCQAGDLRDMISMYLPG
AEVPEPAAPSRLHMSQHYQSGPVPGTAINGTLPLSHM
```

##### 3.3.2. Sequence alignment: vertebrates

```
sox2-human          MYNMMETELKPPGPQQTSGGGGGNSTAAAAGGNQKNSPDRVKRPMNAFMVWSRGQRRKMA 60
sox2-Danio          MYNMMETELKPPAPQPNTGG-TGNTNSSGN--NQKNSPDRIKRPMNAFMVWSRGQRRKMA 57
sox2-Gallus         MYNMMETELKPPAPQQTSGGGTGNSNSAAN--NQKNSPDRVKRPMNAFMVWSRGQRRKMA 58
                    ************.** .:**  **:.::.   ********:*******************

sox2-human          QENPKMHNSEISKRLGAEWKLLSETEKRPFIDEAKRLRALHMKEHPDYKYRPRRKTKTLM 120
sox2-Danio          QENPKMHNSEISKRLGAEWKLLSESEKRPFIDEAKRLRALHMKEHPDYKYRPRRKTKTLM 117
sox2-Gallus         QENPKMHNSEISKRLGAEWKLLSEAEKRPFIDEAKRLRALHMKEHPDYKYRPRRKTKTLM 118
                    ************************:***********************************

sox2-human          KKDKYTLPGGLLAPGGNSMASGVGVGAGLGAGVNQRMDSYAHMNGWSNGSYSMMQDQLGY 180
sox2-Danio          KKDKYTLPGGLLAPGGNGMAGVGVGAGLGAGVNQRMDSYAHMNGWTNGGYGMMQEQLGY 177
```

```
sox2-Gallus      KKDKYTLPGGLLAPGTNTMTTGVGVGATLGAGVNQRMDSYAHMNGWTNGGYGMMQEQLGY 178
                 *************** * * :****** ******************:**.*.***:****

sox2-human       PQHPGLNAHGAAQMQPMHRYDVSALQYNSMTSSQTYMNGSPTYSMSYSQQGTPGMALGSM 240
sox2-Danio       PQHPSLNAHNTAQMQPMHRYDMSALQYNSMTNSQTYMNGSPTYSMSYSQQSTPGMTLGSM 237
sox2-Gallus      PQHPGLNAHNAAQMQPMHRYDVSALQYNSMTSSQTYMNGSPTYSMSYSQQGTPGMALGSM 238
                 ****.****.:**********:*********.*****************.****:****

sox2-human       GSVVKSEASSSPPVVTSSSHSRA-PCQAGDLRDMISMYLPGAEVPEPAAPSRLHMSQHYQ 299
sox2-Danio       GSVVKSESSSSPPVVTSSSHSRAGQCQTGDLRDMISMYLPGAEVQDQSAQSRLHMSQHYQ 297
sox2-Gallus      GSVVKTESSSSPPVVTSSSHSRA-PCQAGDLRDMISMYLPGAEVPEPAAPSRLHMSQHYQ 297
                 *****:*:***************  **:*************** : :* **********

sox2-human       SGPVPGTAINGTLPLSHM 317
sox2-Danio       SAPVPGTTINGTIPLSHM 315
sox2-Gallus      SAPVPGTAINGTLPLSHM 315
                 *.*****:****:*****
```

##### 3.3.3. Disorder prediction

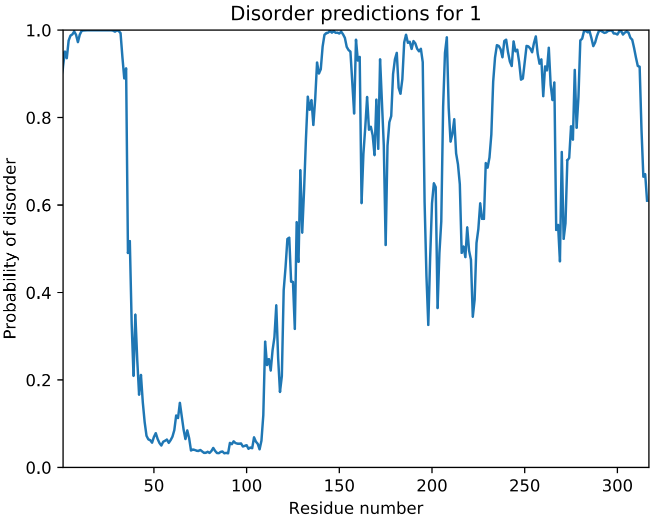


##### 3.3.4. Coding DNA sequences, produced protein constructs

#### Sox2-aa1-42

Synthesized sequence :

ccgcgggtgagaacctgtacttccagggcatgtataacatgatggaaaccgaactgaagccgccgggtccgcagcaaaccagcggtggcggtgg
cggtaacagcaccgctgcggcggcgggtggtaaccaaaagaacagcccggaccgtgtgaaataaaagctt

Translates into :

AGENLYFQGMYNMMETELKPPGPQQTSGGGGGNSTAAAAGGNQKNSPDRVK*

After TEV-cleavage *(leaving 1 Gly in N-ter)*:

```
         10         20         30         40
G MYNMMETEL KPPGPQQTSG GGGGNSTAAA AGGNQKNSPD RVK
```

#### Sox2-aa115-317_C265A

Synthesized sequence :

cgcgggtgagaacctgtacttccagggcaagaccaaaaccctgatgaagaaagacaagtataccctgccgggtggcctgctggcgccgggtggc
aacagcatggcgagcggtgtgggcgttggtgcgggcctgggtgcgggcgtgaaccagcgtatggacagctacgcgcacatgaacggttggagca
acggcagctacagcatgatgcaggatcaactgggttatccgcaacatccgggtctgaacgcgcatggtgcggcgcagatgcaaccgatgcaccg
ttacgatgttagcgcgctgcagtataacagcatgaccagcagccaaacctatatgaacggcagcccgacctacagcatgagctatagccaacaa
ggtaccccgggtatggcgctgggtagcatgggcagcgtggttaaaagcgaagcgagcagcagcccgccggtggttaccagcagcagccacagcc
gtgcgccggcgcaagcgggtgacctgcgtgatatgatcagcatgtacctgccgggtgcggaagtgccggaaccggctgcgccgagccgtctgca
catgagccagcactatcaaagcggtccggttccgggcaccgcgattaacggtaccctgccgctgagccacatgtaaaagctt

Translates into :

AGENLYFQGKTKTLMKKDKYTLPGGLLAPGGNSMASGVGVGAGLGAGVNQRMDSYAHMNGWSNGSYSMMQDQLGYPQHPGLNAHGAAQMQPMHR
YDVSALQYNSMTSSQTYMNGSPTYSMSYSQQGTPGMALGSMGSVVKSEASSSPPVVTSSSHSRAPAQAGDLRDMISMYLPGAEVPEPAAPSRLH
MSQHYQSGPVPGTAINGTLPLSHM*

After TEV-cleavage *(leaving 1 Gly in N-ter)*:

```
     120
G KTKTLM

       130        140        150        160        170        180
KKDKYTLPGG LLAPGGNSMA SGVGVGAGLG AGVNQRMDSY AHMNGWSNGS YSMMQDQLGY

       190        200        210        220        230        240
PQHPGLNAHG AAQMQPMHRY DVSALQYNSM TSSQTYMNGS PTYSMSYSQQ GTPGMALGSM

       250        260        270        280        290        300
GSVVKSEASS SPPVVTSSSH SRAPAQAGDL RDMISMYLPG AEVPEPAAPS RLHMSQHYQS

       310
GPVPGTAING TLPLSHM
```

#### Sox2-aa115-187

Coding sequence – mutation from Sox2-aa115-317 C265A :

Ccgcgggtgagaacctgtacttccagggcaagaccaaaaccctgatgaagaaagacaagtataccctgccgggtggcctgctggcgccgggtgg
caacagcatggcgagcggtgtgggcgttggtgcgggcctgggtgcgggcgtgaaccagcgtatggacagctacgcgcacatgaacggttggagc
aacggcagctacagcatgatgcaggatcaactgggttatccgcaacatccgggtctgaactaaaagctt

Translates into :

AGENLYFQGKTKTLMKKDKYTLPGGLLAPGGNSMASGVGVGAGLGAGVNQRMDSYAHMNGWSNGSYSMMQDQLGYPQHPGLN*

After TEV-cleavage *(leaving 1 Gly in N-ter)*:

```
     120
G KTKTLM

       130        140        150        160        170        180
KKDKYTLPGG LLAPGGNSMA SGVGVGAGLG AGVNQRMDSY AHMNGWSNGS YSMMQDQLGY

       190
PQHPGLN
```

#### Sox2-aa115-236

Coding sequence – mutation from Sox2-aa115-317_C265A :

ccgcgggtgagaacctgtacttccagggcaagaccaaaaccctgatgaagaaagacaagtataccctgccgggtggcctgctggcgccgggtgg
caacagcatggcgagcggtgtgggcgttggtgcgggcctgggtgcgggcgtgaaccagcgtatggacagctacgcgcacatgaacggttggagc
aacggcagctacagcatgatgcaggatcaactgggttatccgcaacatccgggtctgaacgcgcatggtgcggcgcagatgcaaccgatgcacc
gttacgatgttagcgcgctgcagtataacagcatgaccagcagccaaacctatatgaacggcagcccgacctacagcatgagctatagccaaca
aggtaccccgggtatggcgtaaaagctt

Translates into :

AGENLYFQGKTKTLMKKDKYTLPGGLLAPGGNSMASGVGVGAGLGAGVNQRMDSYAHMNGWSNGSYSMMQDQLGYPQHPGLNAHGAAQMQPMHR
YDVSALQYNSMTSSQTYMNGSPTYSMSYSQQGTPGMA*

After TEV-cleavage *(leaving 1 Gly in N-ter)*:

```
     120
G KTKTLM

       130        140        150        160        170        180
KKDKYTLPGG LLAPGGNSMA SGVGVGAGLG AGVNQRMDSY AHMNGWSNGS YSMMQDQLGY

       190        200        210        220        230        240
PQHPGLNAHG AAQMQPMHRY DVSALQYNSM TSSQTYMNGS PTYSMSYSQQ GTPGMA
```

#### Sox2-aa115-282_C265A

Coding sequence – mutation from Sox2-aa115-317_C265A :

ccgcgggtgagaacctgtacttccagggcaagaccaaaaccctgatgaagaaagacaagtataccctgccgggtggcctgctggcgccgggtgg
caacagcatggcgagcggtgtgggcgttggtgcgggcctgggtgcgggcgtgaaccagcgtatggacagctacgcgcacatgaacggttggagc
aacggcagctacagcatgatgcaggatcaactgggttatccgcaacatccgggtctgaacgcgcatggtgcggcgcagatgcaaccgatgcacc
gttacgatgttagcgcgctgcagtataacagcatgaccagcagccaaacctatatgaacggcagcccgacctacagcatgagctatagccaaca
aggtaccccgggtatggcgctgggtagcatgggcagcgtggttaaaagcgaagcgagcagcagcccgccggtggttaccagcagcagccacagc
cgtgcgccggcgcaagcgggtgacctgcgtgatatgatcagcatgtacctgccgggtgcggaataaaagctt

Translates into :

AGENLYFQGKTKTLMKKDKYTLPGGLLAPGGNSMASGVGVGAGLGAGVNQRMDSYAHMNGWSNGSYSMMQDQLGYPQHPGLNAHGAAQMQPMHR
YDVSALQYNSMTSSQTYMNGSPTYSMSYSQQGTPGMALGSMGSVVKSEASSSPPVVTSSSHSRAPAQAGDLRDMISMYLPGAE*

After TEV-cleavage *(leaving 1 Gly in N-ter)*:

```
     120
G KTKTLM

       130        140        150        160        170        180
KKDKYTLPGG LLAPGGNSMA SGVGVGAGLG AGVNQRMDSY AHMNGWSNGS YSMMQDQLGY

       190        200        210        220        230        240
PQHPGLNAHG AAQMQPMHRY DVSALQYNSM TSSQTYMNGS PTYSMSYSQQ GTPGMALGSM

       250        260        270        280
GSVVKSEASS SPPVVTSSSH SRAPAQAGDL RDMISMYLPG AE
```

#### AviTag-Sox2-115-317_C265A

Coding sequence – mutation from Sox2-aa115-317 C265A :

Ggtaccggcctgaacgacattttttgaagcgcagaagatcgagtggcacgagggcgcgggcaagaccaagaccctgatgaagaaggacaagtata
ccctgccgggtggcctgctggcgccgggtggcaacagcatggcgagcggtgtgggcgttggtgcgggcctgggtgcgggcgtgaaccagcgtat
ggacagctacgcgcacatgaacggttggagcaacggcagctacagcatgatgcaggatcaactgggttatccgcaacatccgggtctgaacgcg
catggtgcggcgcagatgcaaccgatgcaccgttacgacgttagcgcgctgcagtataacagcatgaccagcagccaaacctatatgaacggta
gcccgacctacagcatgagctatagccaacagggcaccccgggtatggcgctgggtagcatgggcagcgtggttaaaagcgaggcgagcagcag
cccgccggtggttaccagcagcagccacagccgtgcgccggcgcaggcgggtgacctgcgtgatatgatcagcatgtacctgccgggtgcggaa
gtgccggaaccggcggcgccgagccgtctgcacatgagccaacactatcagagcggtccggttccgggcaccgcgattaacggcaccctgccgc
tgagccatatgtaaaagctt

Translates into :

GTGLNDIFEAQKIEWHEGAGKTKTLMKKDKYTLPGGLLAPGGNSMASGVGVGAGLGAGVNQRMDSYAHMNGWSNGSYSMMQDQLGYPQHPGLNA
HGAAQMQPMHRYDVSALQYNSMTSSQTYMNGSPTYSMSYSQQGTPGMALGSMGSVVKSEASSSPPVVTSSSHSRAPAQAGDLRDMISMYLPGAE
VPEPAAPSRLHMSQHYQSGPVPGTAINGTLPLSHM*

Expressed peptide:

MAHHHHHHVGTGLNDIFEAQKIEWHEGAGKTKTLMKKDKYTLPGGLLAPGGNSMASGVGVGAGLGAGVNQRMDSYAHMNGWSNGSYSMMQDQLG
YPQHPGLNAHGAAQMQPMHRYDVSALQYNSMTSSQTYMNGSPTYSMSYSQQGTPGMALGSMGSVVKSEASSSPPVVTSSSHSRAPAQAGDLRDM
ISMYLPGAEVPEPAAPSRLHMSQHYQSGPVPGTAINGTLPLSHM*

```
                                     120
MAHHHHHHVGTGLNDIFEAQKIEWHEGAGG KTKTLM

       130        140        150        160        170        180
KKDKYTLPGG LLAPGGNSMA SGVGVGAGLG AGVNQRMDSY AHMNGWSNGS YSMMQDQLGY

       190        200        210        220        230        240
PQHPGLNAHG AAQMQPMHRY DVSALQYNSM TSSQTYMNGS PTYSMSYSQQ GTPGMALGSM

       250        260        270        280        290        300
GSVVKSEASS SPPVVTSSSH SRAPAQAGDL RDMISMYLPG AEVPEPAAPS RLHMSQHYQS

       310
GPVPGTAING TLPLSHM
```

#### AviTag-Sox2-115-240

Coding sequence – mutation from Sox2-aa115-317 C265A :

Ggtaccggcctgaacgacattttttgaagcgcagaagatcgagtggcacgagggcgcgggcaagaccaagaccctgatgaagaaggacaagtata
ccctgccgggtggcctgctggcgccgggtggcaacagcatggcgagcggtgtgggcgttggtgcgggcctgggtgcgggcgtgaaccagcgtat
ggacagctacgcgcacatgaacggttggagcaacggcagctacagcatgatgcaggatcaactgggttatccgcaacatccgggtctgaacgcg
catggtgcggcgcagatgcaaccgatgcaccgttacgacgttagcgcgctgcagtataacagcatgaccagcagccaaacctatatgaacggta
gcccgacctacagcatgagctatagccaacagggcaccccgggtatggcgctgggtagcatgtaaaagctt

Translates into :

GTGLNDIFEAQKIEWHEGAGKTKTLMKKDKYTLPGGLLAPGGNSMASGVGVGAGLGAGVNQRMDSYAHMNGWSNGSYSMMQDQLGYPQHPGLNA
HGAAQMQPMHRYDVSALQYNSMTSSQTYMNGSPTYSMSYSQQGTPGMALGSM*

Expressed peptide:

MAHHHHHHVGTGLNDIFEAQKIEWHEGAGKTKTLMKKDKYTLPGGLLAPGGNSMASGVGVGAGLGAGVNQRMDSYAHMNGWSNGSYSMMQDQLG
YPQHPGLNAHGAAQMQPMHRYDVSALQYNSMTSSQTYMNGSPTYSMSYSQQGTPGMALGSM*

```
                                     120
MAHHHHHHVGTGLNDIFEAQKIEWHEGAGG KTKTLM

       130        140        150        160        170        180
KKDKYTLPGG LLAPGGNSMA SGVGVGAGLG AGVNQRMDSY AHMNGWSNGS YSMMQDQLGY

       190        200        210        220        230        240
PQHPGLNAHG AAQMQPMHRY DVSALQYNSM TSSQTYMNGS PTYSMSYSQQ GTPGMALGSM
```

#### AviTag-Sox2-234-317_C265A

Coding sequence – mutation from Sox2-aa115-317 C265A :

```
ggtaccggcctgaacgacatttttgaagcgcagaagatcgagtggcacgagggcgcgggtatggcgctgggtagcatgggcagcgtggttaaaa
gcgaggcgagcagcagcccgccggtggttaccagcagcagccacagccgtgcgccggcgcaggcgggtgacctgcgtgatatgatcagcatgta
cctgccgggtgcggaagtgccggaaccggcggcgccgagccgtctgcacatgagccaacactatcagagcggtccggttccgggcaccgcgatt
aacggcaccctgccgctgagccatatgtaaaagctt
```

Translates into :

```
GTGLNDIFEAQKIEWHEGAGMALGSMGSVVKSEASSSPPVVTSSSHSRAPAQAGDLRDMISMYLPGAEVPEPAAPSRLHMSQHYQSGPVPGTAI
NGTLPLSHM*
```

Expressed peptide:

```
MAHHHHHHVGTGLNDIFEAQKIEWHEGAGMALGSMGSVVKSEASSSPPVVTSSSHSRAPAQAGDLRDMISMYLPGAEVPEPAAPSRLHMSQHYQ
SGPVPGTAINGTLPLSHM*
```

```
                                    140
MAHHHHHHVGTGLNDIFEAQKIEWHEGAG MALGSM

       250        260        270        280        290        300
GSVVKSEASS SPPVVTSSSH SRAPAQAGDL RDMISMYLPG AEVPEPAAPS RLHMSQHYQS

       310
GPVPGTAING TLPLSHM
```

#### 3.4. Nanog

##### 3.4.1. Sequence alignment: mammals

```
S: Serine
P : Proline
SP : phosphorylation motif for MAPKand Cdk
FILVYW : hydrophobic
DE : Asp/Glu
KR : Lys/Arg
T : Thr
XXX: DND-BD in crystal (4RBO)

Nanog-human             MSVDPACPQSLP-CFEASDCKESSPMPVICGPEENYPSLQMSSAEMPHTETVSPLPS-SM 58
nanog-Mus               MSVGLPGPHSLPSSEEASNSGNASSMPAVFHP-ENYSCLQGSATEMLCTEAASPRPS-SE 58
NANOG-Bos               MSVGPACPQSLL-GPEASNSRESSPMP-----EESYVSLQTSSADTLDTDTVSPLPS-SM 53
Nanog-Canis             ---------MPA-GPQAPNSRDPSPMPEVYGPRGNPASLPMSSAETPHAETVSPLPS-SM 49
Nanog-Capra             MSVDPACPQSLL-GPEASNSGESSPMP-----EESYASLQMSSADTLDTDTVSPLPS-SM 53
NANOG-Balaenoptera      MSVDPACPQSLR-GPEASNSRESSPMPEIYGPEENYVSLQMSSVETHDMETVSPLPSFSM 59
                                       :*. . :.*.:*       .   *  *:.:    ::.** ** .

Nanog-human             DLLIQDSPDSSTSPKGK-QPTSAEK-SVAKKEDKVPVKKQKTRTVFSSTQLCVLNDRFQR 116
nanog-Mus               DLPLQGSPDSSTSPKQKLSSPEADKGPEEEE-NKVLARKQKMRTVFSQAQLCALKDRFQK 117
NANOG-Bos               DLLIQDSPDSSTSPRVKPLSPSVEE-STEK-EETVPVKKQKIRTVFSQTQLCVLNDRFQR 111
Nanog-Canis             DLLTQDSPDSSTSPRVKLPPTSGEE-RTARKEDATQGKKQKMRTVFSQTQLYVLNDRFQR 108
Nanog-Capra             DLLIHDNPDSSTSPRVKPLSPSAEE-STEK-EEKVPVKKQKIRTVFSQTQLCVLNDRFQR 111
NANOG-Balaenoptera      DLLIQDSPDSSTSPRVKLLATAADK-STEKKEEKVLIKKQKTRTVFSQTQLCVLNDRFQR 118
                        **  :..****:.*: *   .   .    .       :*** *****.:** .*:****:

Nanog-human             QKYLSLQQMQELSNILNLSYKQVKTWFQNQRMKSKRWQKNNWPKNSNGVTQKA-SAPTYP 175
nanog-Mus               QKYLSLQQMQELSSILNLSYKQVKTWFQNQRMKCKRWQKNQWLKTSNGLIQKGSAPVEYP 177
NANOG-Bos               QKYLSLQQMQELSNILNLSYKQVKTWFQNQRMKCKKWQKNNWPRNSNGMPQGP-AMAEYP 170
Nanog-Canis             QKYLSLQQMQELSNILNLSYKQVKTWFQNQRMKSKRWQKSNWPKESNSVTQNSSATTEYA 168
Nanog-Capra             QKYLSLQQMQELSNILNLSYKQVKTWFQNQRMKCKKWQKNNWPRNSNDVPQDP-ATAEYP 170
NANOG-Balaenoptera      QKYLSLQQMQELSNILNLSYKQVKTWFQNQRMKCKRWQKNNWPRNSNTVTQGP-ATTEYP 177
                        *:********:**.**.***************.*:***.:*  : .* : *   :   *.

Nanog-human             SLYSSYHQGCLVNPTGNLPMWS----------NQTWNNSTWSNQTQNIQSWSNHSWNTQT 225
nanog-Mus               SIHCSYPQGYLVNASGSLSMWGSQTWTNPTWSSQTWTNPTWNNQTWTNPTWSSQAWTAQS 237
NANOG-Bos               GFYS-YHQGCLVNSPGNLPMWG----------NQTWNNPTWSNQSWNSQSWSNHSWNSQA 219
Nanog-Canis             GFYP-CRQGYLLNPSGNLPLW----------SSQAWNNPNWSSQTWNSQSWSSHSWNSQT 217
Nanog-Capra             SFYS-YHQGCLVNSPRNMPMWG----------NQTWNNPTWSNQNWNSQSWSNHSWNSQA 219
NANOG-Balaenoptera      GFYS-YHQGCLANSSGNLPMWG----------NQTWNNPTWSNQSWNSQSWSNHPWNNQT 226
                        .::    ** * *.. .:.:*           .*:*.. .*..*. .  :**.:.*. *:

Nanog-human             WCTQSWNNQAWNSP-FYNCGEESLQSCMQFQPNSPASDLEAALEAAGEGLNVIQQTTRYF 284
nanog-Mus               W-----NGQPWNAAPLHNFGEDFLQPYVQLQQNFSASDLEVNLEATRESH-------AHF 285
NANOG-Bos               WCPQAWNNQPWNNQ-FNNYMEEFLQPGIQLQQNSPVCDLEATLGTAGENYNVIQQTVKYF 278
Nanog-Canis             WCPQAWNNQAWNNP-LHNCEEESLQPPIQFQQNS-MGDLESIFETAGESHGVLQQSTKYF 275
Nanog-Capra             WCPQAWNNQPWNNQ-CNNYMEEFLQPGIQLQQNSPVCDLEATLGTAGENYNVIQQAVKYF 278
NANOG-Balaenoptera      WCPQAWNNQTSNNQ-FNNYVEEFLQPQTQFQQNSPVSDLEATLETAGESYNIIQQTAKYF 285
                       *     *.*  *     :  *:  *.   :* *    **:  : :: *.         :

Nanog-human             STP-QTMDLFLNYSMNMQPEDV 305
nanog-Mus               STP-QALELFLNYSVTP-PGEI 305
NANOG-Bos               NSQQQITDLFPNYPLNIQPEDL 300
Nanog-Canis             STP-QIMDFFPNYSXNIQPEDV 296
Nanog-Capra             SSQQQITDLFPNYPLNIQPEDL 300
NANOG-Balaenoptera      NSQQQIMDLFPNYSLNIQPEDL 307
                        .:  *  ::* **.
```

```
>Nanog-human
MSVDPACPQSLPCFEASDCKESSPMPVICGPEENYPSLQMSSAEMPHTETVSPLPSSMDLLIQDSPDSST
SPKGKQPTSAEKSVAKKEDKVPVKKQKTRTVFSSTQLCVLNDRFQRQKYLSLQQMQELSNILNLSYKQVK
TWFQNQRMKSKRWQKNNWPKNSNGVTQKASAPTYPSLYSSYHQGCLVNPTGNLPMWSNQTWNNSTWSNQT
QNIQSWSNHSWNTQTWCTQSWNNQAWNSPFYNCGEESLQSCMQFQPNSPASDLEAALEAAGEGLNVIQQT
TRYFSTPQTMDLFLNYSMNMQPEDV
```

##### 3.4.2. Sequence alignment: vertebrates

##### 3.4.3. Disorder prediction

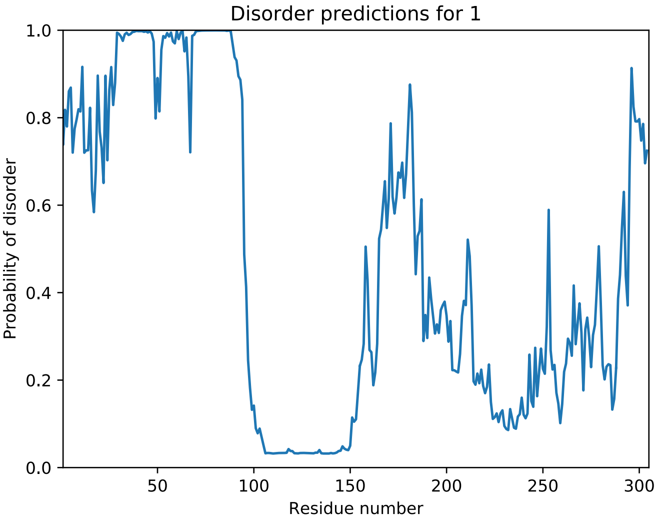


#### 3.4.4. Coding DNA sequences, produced protein constructs

##### Nanog-aa1-85

Name Genscript : TEV-NanogNter

Coding sequence (mutated from Nanog-aa1-85-Tev):

Caattggtgaaaatctgtacttccagggcatgtccgtcgatccggcgtgtccgcagagcctgccgtgctttgaagcgagcgactgtaaagaatc
gagcccgatgccggtcatttgcggcccggaagaaaactatccgtctctgcagatgagctctgcagaaatgccgcatacggaaaccgtgagcccg
ctgccgagttccatggatctgctgatccaggatagtccggactcatcgacgtccccgaaaggtaaacaaccgaccagcgcggaaaaatctgtgg
cctaaaagctt

Translates into :

IGENLYFQGMSVDPACPQSLPCFEASDCKESSPMPVICGPEENYPSLQMSSAEMPHTETVSPLPSSMDLLIQDSPDSSTSPKGKQPTSAEKSVA
*

Before TEV-cleavage:

GST-His-Tev_NanogCter_aa1-85

MSPILGYWKIKGLVQPTRLLLEYLEEKYEEHLYERDEGDKWRNKKFELGLEFPNLPYYIDGDVKLTQSMAIIRYIADKHNMLGGCPKERAEISM
LEGAVLDIRYGVSRIAYSKDFETLKVDFLSKLPEMLKMFEDRLCHKTYLNGDHVTHPDFMLYDALDVVLYMDPMCLDAFPKLVCFKKRIEAIPQ
IDKYLKSSKYIAWPLQGWQATFGGGDHPPKSDGSTSGSGHHHHHHSAGLVPRGSTAIGENLYFQGMSVDPACPQSLPCFEASDCKESSPMPVIC
GPEENYPSLQMSSAEMPHTETVSPLPSSMDLLIQDSPDSSTSPKGKQPTSAEKSVA*
Number of amino acids: 338
Molecular weight: 38016.43
Theoretical pI: 5.54
Total number of negatively charged residues (Asp + Glu): 47
Total number of positively charged residues (Arg + Lys): 35
Ext. coefficient 46340
Abs 0.1% (=1 g/l) 1.219, assuming all pairs of Cys residues form cystines
Ext. coefficient 45840
Abs 0.1% (=1 g/l) 1.206, assuming all Cys residues are reduced

After Tev-cleavage *(leaving 1 Gly in N-ter)*:

GMSVDPACPQSLPCFEASDCKESSPMPVICGPEENYPSLQMSSAEMPHTETVSPLPSSMDLLIQDSPDSSTSPKGKQPTSAEKSVA

```
          10         20         30         40         50         60
G MSVDPACPQS LPCFEASDCK ESSPMPVICG PEENYPSLQM SSAEMPHTET VSPLPSSMDL

        70         80
LIQDSPDSST SPKGKQPTSA EKSVA
```

##### Nanog-aa154-305-Tev

Synthesized sequence :

Ccgcgggtgagaacctgtacttccaaggcatggcgggtcacctggcgagcgattttgcgtttagcccgccgccgggtggtggtggtgacggtcc
gggtggcccggaaccgggttgggtggatccgcgtacctggctgagcttccaaggtccgccgggtggcccgggtattggtccgggtgtgggcccg
ggtagcgaggtttggggtattccgccgtgcccgccgccgtacgaattttgcggtggcatggcgtattgcggtccgcaagtgggcgttggtctgg
ttccgcaaggtggcctggaaaccagccagccggagggtgaagcgggcgtgggtgttgagagcaacagcgatggtgcgagcccggaaccgtgcac
cgtgaccccgggtgcggttaagctggagaaggaaaaactggagcagaacccggaggaaagccaagatatcaaggcgctgcagaaagaaaacctg

tactttcaaggtggcgcgggtggcgcgggtggccagtataaactgattctgaacggcaagaccctgaaaggtgaaaccaccaccgaagcggtgg
atgcggcgaccgctgagaaggttttcaaacagtacgcgaacgacaacggcgtggatggcgagtggacctatgacgatgcgaccaagacctttac
cgttaccgaaggtggctaaaagctt

Translates into :
ENLYFQGQKNNWPKNSNGVTQKASAPTYPSLYSSYHQGCLVNPTGNLPMWSNQTWNNSTWSNQTQNIQSWSNHSWNTQTWCTQSWNNQAWNSPF
YNCGEESLQSCMQFQPNSPASDLEAALEAAGEGLNVIQQTTRYFSTPQTMDLFLNYSMNMQPEDVENLYFQGGAGGAGGQYKLILNGKTLKGET
TTEAVDAATAEKVFKQYANDNGVDGEWTYDDATKTFTVTEGG*

Bfore TEV-cleavage:

GST-His-Tev_NanogCter_aa154-305_Tev-GB1 from pET41a+

MSPILGYWKIKGLVQPTRLLLEYLEEKYEEHLYERDEGDKWRNKKFELGLEFPNLPYYIDGDVKLTQSMAIIRYIADKHNMLGGCPKERAEISM
LEGAVLDIRYGVSRIAYSKDFETLKVDFLSKLPEMLKMFEDRLCHKTYLNGDHVTHPDFMLYDALDVVLYMDPMCLDAFPKLVCFKKRIEAIPQ
IDKYLKSSKYIAWPLQGWQATFGGGDHPPKSDGSTSGSGHHHHHHSAGLVPRGSTAIGMKETAENLYFQGQKNNWPKNSNGVTQKASAPTYPSL
YSSYHQGCLVNPTGNLPMWSNQTWNNSTWSNQTQNIQSWSNHSWNTQTWCTQSWNNQAWNSPFYNCGEESLQSCMQFQPNSPASDLEAALEAAG
EGLNVIQQTTRYFSTPQTMDLFLNYSMNMQPEDVENLYFQGGAGGAGGQYKLILNGKTLKGETTTEAVDAATAEKVFKQYANDNGVDGEWTYDD
ATKTFTVTEGG

After Tev-cleavage *(leaving 1 Gly in N-ter and ENLYFQ in C-ter)*:

GQKNNWPKNSNGVTQKASAPTYPSLYSSYHQGCLVNPTGNLPMWSNQTWNNSTWSNQTQNIQSWSNHSWNTQTWCTQSWNNQAWNSPFYNCGEE
SLQSCMQFQPNSPASDLEAALEAAGEGLNVIQQTTRYFSTPQTMDLFLNYSMNMQPEDVENLYFQ

```
      160        170        180
G QKNNWPK NSNGVTQKAS APTYPSLYSS

       190        200        210        220        230        240
YHQGCLVNPT GNLPMWSNQT WNNSTWSNQT QNIQSWSNHS WNTQTWCTQS WNNQAWNSPF

       250        260        270        280        290        300
YNCGEESLQS CMQFQPNSPA SDLEAALEAA GEGLNVIQQT TRYFSTPQTM DLFLNYSMNM


QPEDV ENLYFQ
```

#### Nanog-aa154-305

Coding sequence (mutated from Nanog-aa154-305-Tev):
ccgcgggtgaaaacctgtacttccagggtcaaaagaacaactggccgaaaaacagcaacggtgtgacccaaaaggcgagcgcgccgacctatcc
gagcctgtacagcagctatcaccagggttgcctggttaacccgaccggcaacctgccgatgtggagcaaccaaacctggaacaacagcacctgg
agcaaccagacccaaaacatccagagctggagcaaccacagctggaacacccagacctggtgcacccaaagctggaacaaccaggcgtggaaca
gcccgttctacaactgcggcgaggaaagcctgcaaagctgcatgcagtttcaaccgaacagcccggcgagcgacctggaggcggcgctggaagc
ggcgggtgaaggcctgaacgtgatccagcaaaccacccgttacttcagcaccccgcaaaccatggacctgtttctgaactatagcatgaacatg
cagccggaggatgtttaaaagctt

Translates into :
ENLYFQGQKNNWPKNSNGVTQKASAPTYPSLYSSYHQGCLVNPTGNLPMWSNQTWNNSTWSNQTQNIQSWSNHSWNTQTWCTQSWNNQAWNSPF
YNCGEESLQSCMQFQPNSPASDLEAALEAAGEGLNVIQQTTRYFSTPQTMDLFLNYSMNMQPEDVENLYFQGGAGGAGGQYKLILNGKTLKGET
TTEAVDAATAEKVFKQYANDNGVDGEWTYDDATKTFTVTEGG*

After Tev-cleavage *(leaving 1 Gly in N-ter):*

GQKNNWPKNSNGVTQKASAPTYPSLYSSYHQGCLVNPTGNLPMWSNQTWNNSTWSNQTQNIQSWSNHSWNTQTWCTQSWNNQAWNSPFYNCGEE
SLQSCMQFQPNSPASDLEAALEAAGEGLNVIQQTTRYFSTPQTMDLFLNYSMNMQPEDV*

```
      160        170        180
G QKNNWPK NSNGVTQKAS APTYPSLYSS

       190        200        210        220        230        240
YHQGCLVNPT GNLPMWSNQT WNNSTWSNQT QNIQSWSNHS WNTQTWCTQS WNNQAWNSPF

       250        260        270        280        290        300
YNCGEESLQS CMQFQPNSPA SDLEAALEAA GEGLNVIQQT TRYFSTPQTM DLFLNYSMNM


QPEDV
```

#### Nanog-aa154-215-Tev

Coding sequence (mutated from Nanog-aa154-305-Tev):
Ccgcgggtgaaaacctgtacttccagggtcaaaagaacaactggccgaaaaacagcaacggtgtgacccaaaaggcgagcgcgccgacctatcc
gagcctgtacagcagctatcaccagggttgcctggttaacccgaccggcaacctgccgatgtggagcaaccaaacctggaacaacagcacctgg
agcaaccagacccaaaacatccagagcgaaaacctgtactttcaaggtggcgcgggtggcgcgggtggccagtataagctgattctgaacggca
agaccctgaaaggcgaaaccaccaccgaggcggtggatgcggcgaccgctgagaaggttttcaaacagtacgcgaacgacaacggtgtggatgg
cgaatggacctatgacgatgcgaccaaaacctttaccgttaccgagggtggctaaaagctt

<u>Translates into:</u>
AGENLYFQGQKNNWPKNSNGVTQKASAPTYPSLYSSYHQGCLVNPTGNLPMWSNQTWNNSTWSNQTQNIQSENLYFQGGAGGAGGQYKLILNGK
TLKGETTTEAVDAATAEKVFKQYANDNGVDGEWTYDDATKTFTVTEGG

<u>After Tev-cleavage *(leaving 1 Gly in N-ter and ENLYFQ in C-ter):*</u>

GQKNNWPKNSNGVTQKASAPTYPSLYSSYHQGCLVNPTGNLPMWSNQTWNNSTWSNQTQNIQSENLYFQ

```
      160        170        180
G QKNNWPK NSNGVTQKAS APTYPSLYSS

       190        200        210
YHQGCLVNPT GNLPMWSNQT WNNSTWSNQT QNIQS ENLYFQ
```

#### Nanog-aa154-272-Tev

<u>Coding sequence (mutated from Nanog-aa154-305-Tev):</u>
Ccgcgggtgaaaacctgtacttccagggtcaaaagaacaactggccgaaaaacagcaacggtgtgacccaaaaggcgagcgcgccgacctatcc
gagcctgtacagcagctatcaccagggttgcctggttaacccgaccggcaacctgccgatgtggagcaaccaaacctggaacaacagcacctgg
agcaaccagacccaaaacatccagagctggagcaaccacagctggaacacccagacctggtgcacccaaagctggaacaaccaggcgtggaaca
gcccgttctacaactgcggcgaggaaagcctgcaaagctgcatgcagtttcaaccgaacagcccggcgagcgacctggaggcggcgctggaagc
ggcgggtgaagaaaacctgtactttcaaggtggcgcgggtggcgcgggtggccagtataagctgattctgaacggcaagaccctgaaaggcgaa
accaccaccgaggcggtggatgcggcgaccgctgagaaggttttcaaacagtacgcgaacgacaacggtgtggatggcgaatggacctatgacg
atgcgaccaaaacctttaccgttaccgagggtggctaaaagctt

<u>Translates into:</u>
AGENLYFQGQKNNWPKNSNGVTQKASAPTYPSLYSSYHQGCLVNPTGNLPMWSNQTWNNSTWSNQTQNIQSWSNHSWNTQTWCTQSWNNQAWNS
PFYNCGEESLQSCMQFQPNSPASDLEAALEAAGEENLYFQGGAGGAGGQYKLILNGKTLKGETTTEAVDAATAEKVFKQYANDNGVDGEWTYDD
ATKTFTVTEGG*

<u>After Tev-cleavage *(leaving 1 Gly in N-ter and ENLYFQ in C-ter):*</u>

GQKNNWPKNSNGVTQKASAPTYPSLYSSYHQGCLVNPTGNLPMWSNQTWNNSTWSNQTQNIQSWSNHSWNTQTWCTQSWNNQAWNSPFYNCGEE
SLQSCMQFQPNSPASDLEAALEAAGEENLYFQ

```
      160        170        180
G QKNNWPK NSNGVTQKAS APTYPSLYSS

       190        200        210        220        230        240
YHQGCLVNPT GNLPMWSNQT WNNSTWSNQT QNIQSWSNHS WNTQTWCTQS WNNQAWNSPF

       250        260        270
YNCGEESLQS CMQFQPNSPA SDLEAALEAA GE ENLYFQG
```

#### Nanog-aa154-305_4C4A

<u>Coding sequence (mutated from Nanog-aa154-305-Tev):</u>
Ccgcgggtgagaatctgtatttccaaggcaaaaacaactggccgaagaacagcaatggtgtgacccaaaaagcgagcgcgccgacctatccgag
cctgtacagcagctatcaccaaggtgcgctggtgaacccgaccggtaacctgccgatgtggagcaaccaaacctggaacaacagcacctggagc
aaccagacccaaaacatccagagctggagcaaccacagctggaacacccagacctgggcgacccaaagctggaacaaccaggcgtggaacagcc
cgttctacaacgcgggcgaggaaagcctgcagagcgcgatgcagtttcaaccgaacagcccggcgagcgacctggaggcggcgctggaagcggc
gggtgaaggcctgaacgttattcagcaaaccacccgttatttcagcaccccgcaaaccatggacctgtttctgaattatagcatgaatatgcag
ccggaggatgtgtaaaagctt

<u>Translates into:</u>
AGENLYFQGKNNWPKNSNGVTQKASAPTYPSLYSSYHQGALVNPTGNLPMWSNQTWNNSTWSNQTQNIQSWSNHSWNTQTWATQSWNNQAWNSP
FYNAGEESLQSAMQFQPNSPASDLEAALEAAGEGLNVIQQTTRYFSTPQTMDLFLNYSMNMQPEDV*

<u>After Tev-cleavage *(leaving 1 Gly in N-ter):*</u>

GKNNWPKNSNGVTQKASAPTYPSLYSSYHQGALVNPTGNLPMWSNQTWNNSTWSNQTQNIQSWSNHSWNTQTWATQSWNNQAWNSPFYNAGEES
LQSAMQFQPNSPASDLEAALEAAGEGLNVIQQTTRYFSTPQTMDLFLNYSMNMQPEDV

```
      160        170        180
G QKNNWPK NSNGVTQKAS APTYPSLYSS

       190        200        210        220        230        240
YHQGALVNPT GNLPMWSNQT WNNSTWSNQT QNIQSWSNHS WNTQTWATQS WNNQAWNSPF

       250        260        270        280        290        300
YNAGEESLQS AMQFQPNSPA SDLEAALEAA GEGLNVIQQT TRYFSTPQTM DLFLNYSMNM


QPEDV
```

#### Nanog-aa154-272_4C4A

Coding sequence (mutated from Nanog-aa154-305 4C4A):

```
Ccgcgggtgagaatctgtatttccaaggcaaaaacaactggccgaagaacagcaatggtgtgacccaaaaagcgagcgcgccgacctatccgag
cctgtacagcagctatcaccaaggtgcgctggtgaacccgaccggtaacctgccgatgtggagcaaccaaacctggaacaacagcacctggagc
aaccagacccaaaacatccagagctggagcaaccacagctggaacacccagacctgggcgacccaaagctggaacaaccaggcgtggaacagcc
cgttctacaacgcgggcgaggaaagcctgcagagcgcgatgcagtttcaaccgaacagcccggcgagcgacctggaggcggcgctggaagcggc
gggtgaataaaagctt
```

Translates into:

```
AGENLYFQGKNNWPKNSNGVTQKASAPTYPSLYSSYHQGALVNPTGNLPMWSNQTWNNSTWSNQTQNIQSWSNHSWNTQTWATQSWNNQAWNSP
FYNAGEESLQSAMQFQPNSPASDLEAALEAAGE
```

After Tev-cleavage *(leaving 1 Gly in N-ter):*

```
GKNNWPKNSNGVTQKASAPTYPSLYSSYHQGALVNPTGNLPMWSNQTWNNSTWSNQTQNIQSWSNHSWNTQTWATQSWNNQAWNSPFYNAGEES
LQSAMQFQPNSPASDLEAALEAAGE

      160        170        180
G QKNNWPK NSNGVTQKAS APTYPSLYSS

       190        200        210        220        230        240
YHQGALVNPT GNLPMWSNQT WNNSTWSNQT QNIQSWSNHS WNTQTWATQS WNNQAWNSPF

       250        260        270
YNAGEESLQS AMQFQPNSPA SDLEAALEAA GE
```

#### 3.5. Esrrb

##### 3.5.1. Sequence alignment: mammals

S: Serine
P : Proline
**SP : phosphorylation motif for MAPKand Cdk**
FILVYW : hydrophobic
DE : Asp/Glu
KR : Lys/Arg
T : Thr
C : Cys

```
human          ---------------------MSSEDRHLGSSCGSFIKTEPSSPSSGIDALSHHSPSGSS
mouse          ---------------------MSSEDRHLGSSCGSFIKTEPSSPSSGIDALSHHSPSGSS
capra          MDVSELCVPDPLGYHNQLLNRMSADDRHLSSSCGSFIKTEPSSPSSGIDALSHHSPSGSS
bos            MDVSELCVPDPLGYHNQLLNRMSADDRHLGSSCGSFIKTEPSSPSSGIDALSHHSPSGSS
balaenoptera   MDVSELCIPDPLGYHNQLLNRMSADDRHLVSSCGSFIKTEPSSPSSGIDALSHHSPRGSS
                                    **::**** ************************** ***

human          DASGGFGIALSTHANGLDSPPMFAGAGLGGNPCRKSYEDCTSGIMEDSAIKCEYMLNAIP
mouse          DASGGFGIALSTHANGLDSPPMFAGAGLGGNPCRKSYEDCTSGIMEDSAIKCEYMLNAIP
capra          DASGGFGLALGAHANGLDSPPMFAGAGLGGTPCRKGYEDCAGGLMEDSAIKCEYMLNAIP
bos            DASGGFGLALGAHANGLDSPPMFAGAGLGGTPCRKGYEDCAGGLMEDSAIKCEYMLNAIP
balaenoptera   DASGGFGLALGAHANGLDSPPMFAGAGLGGTPCRKGYEDCASGIMEDSAIKCEYMLNAIP
               *******:**.:******************.****.****:.*:****************

human          KRL
mouse          KRL
capra          KRL
bos            KRL
balaenoptera   KRL
               ***
```

```
>sp|O95718|ERR2_HUMAN Steroid hormone receptor ERR2 OS=Homo sapiens GN=ESRRB PE=1 SV=2
MSSDDRHLGSSCGSFIKTEPSSPSSGIDALSHHSPSGSSDASGGFGLALGTHANGLDSPP
MFAGAGLGGTPCRKSYEDCASGIMEDSAIKCEYMLNAIPKRLCLVCGDIASGYHYGVASC
EACKAFFKRTIQGNIEYSCPATNECEITKRRRKSCQACRFMKCLKVGMLKEGVRLDRVRG
GRQKYKRRLDSESSPYLSLQISPPAKKPLTKIVSYLLVAEPDKLYAMPPPGMPEGDIKAL
TTLCDLADRELVVIIGWAKHIPGFSSLSLGDQMSLLQSAWMEILILGIVYRSLPYDDKLV
YAEDYIMDEEHSRLAGLLELYRAILQLVRRYKKLKVEKEEFVTLKALALANSDSMYIEDL
EAVQKLQDLLHEALQDYELSQRHEEPWRTGKLLLTLPLLRQTAAKAVQHFYSVKLQGKVP
MHKLFLEMLEAKVGQEQLRGSPKDERMSSHDGKCPFQSAAFTSRDQSNSPGIPNPRPSSP TPLNERGRQISPSTRTPGGQGKHLWLTM
```

##### 3.5.2. Sequence alignment: vertebrates

C: cysteines in N-ter
XX: DNA-BD ordered in NMR structure 1LO1 (construct: human aa96-194_C163A)
*XX*: folded: aa235-432 in Ligand-Binding Domain crystal structures 6LIT, 6LN4 (construct: human aa204-433_Y215H, mutation for solubility/stability)

```
mouse      ---------------------MSSEDRHLGSSCGSFIKTEPSSPSSGIDALSHHSPSGSS   39
human      ---------------------MSSDDRHLGSSCGSFIKTEPSSPSSGIDALSHHSPSGSS   39
goat       MDVSELCVPDPLGYHNQLLNRMSADDRHLSSSCGSFIKTEPSSPSSGIDALSHHSPSGSS   60
whale      MDVSELCIPDPLGYHNQLLNRMSADDRHLVSSCGSFIKTEPSSPSSGIDALSHHSPRGSS   60
chicken    MDISELCISDPLGYHNQLLGRMATEERHLSSSCGSFIKTEPSSPSSGIDAISHHSPSGSS   60
fish       ---------------------MAADERHLPSSCGSYIKTEPSSPSSVIDTVSHHSPSGNS   39
                                *::::*** *****:********** **::***** *.*

mouse      DASGGFGIALSTHANGLDSPPMFAGAGLGGNPCRKSYEDCTSGIMEDSAIKCEYMLNAIP   99
human      DASGGFGLALGTHANGLDSPPMFAGAGLGGTPCRKSYEDCASGIMEDSAIKCEYMLNAIP   99
goat       DASGGFGLALGAHANGLDSPPMFAGAGLGGTPCRKGYEDCAGGLMEDSAIKCEYMLNAIP   120
whale      DASGGFGLALGAHANGLDSPPMFAGAGLGGTPCRKGYEDCASGIMEDSAIKCEYMLNAIP   120
chicken    DASGGYGIAMGGHPNGLDSPPMFNGTGIGGGSCRKRYDDCASAIMEDSPTKCEYMLNAIP   120
fish       DASGGYVSTMNSHSNGLDSPPMFTPSGLGAGTCRKRYDDCSSTIMEDSSIKCEYMLNSLP   99
           *****:  ::. * *********  :*:*.  *** *:**:. :****  *******::*

mouse      KRLCLVCGDIASGYHYGVASCEACKAFFKRTIQGNIEYNCPATNECEITKRRRKSCQACR   159
human      KRLCLVCGDIASGYHYGVASCEACKAFFKRTIQGNIEYSCPATNECEITKRRRKSCQACR   159
goat       KRLCLVCGDIASGYHYGVASCEACKAFFKRTIQGNIEYSCPATNECEITKRRRKSCQACR   180
whale      KRLCLVCGDIASGYHYGVASCEACKAFFKRTIQGNIEYSCPATNECEITKRRRKSCQACR   180
chicken    KRLCLVCGDIASGYHYGVASCEACKAFFKRTIQGNIEYSCPATNECEITKRRRKSCQACR   180
fish       KRLCLVCGDIASGYHYGVASCEACKAFFKRTIQGNIEYSCPATNECEITKRRRKSCQACR   159
           **************************************.*********************

mouse      FMKCLKVGMLKEGVRLDRVRGGRQKYKRRLDSENSPYLNLPISPPAKKPLTKIVSNLLGV   219
human      FMKCLKVGMLKEGVRLDRVRGGRQKYKRRLDSESSPYLSLQISPPAKKPLTKIVSYLLVA   219
goat       FMKCLKVGMLKEGVRLDRVRGGRQKYKRRLDSESSPYLSLQISPPAKKPLTKIVSYLLVA   240
whale      FMKCLKVGMLKEGVRLDRVRGGRQKYKRRLDSESSPYLSLQISPPAKKPLTKIVSYLLVA   240
chicken    FMKCLKVGMLKEGVRLDRVRGGRQKYKRRLDSESSTYLSLQIPPPAKKPLTKIVSHLLVA   240
fish       FMKCLKVGMLKEGVRLDRVRGGRQKYKRRLDSENNPYLGLTLPPPTKKPLTKIVSHLLVA   219
           *********************************.. **.* : **:********* ** .
```

```
mouse        EQDKLYAMPPNDIPEGDIKALTTLCELADRELVFLINWAKHIPGFPSLTLGDQMSLLQSA   279
human        EPDKLYAMPPPGMPEGDIKALTTLCDLADRELVVIIGWAKHIPGFSSLSLGDQMSLLQSA   279
goat         EPDKLYAMPPPGMPEGDIKALTTLCDLADRELVVIIGWAKHIPGFSNLSLGDQMSLLQSA   300
whale        EPDKLYAMPPPGMPEGDIKALTTLCDLADRELVVIIGWAKHIPGFSNLSLGDQMSLLQSA   300
chicken      EPEKIYAMPDPTMPESDIKALTTLCDLADRELVVIIGWAKHIPGFSNLSLGDQMSLLQSA   300
fish         EPEKIYAMPDPTMPESDIKALTTLCDLADRELVVIIGWAKHIPGFSTLSLGDQMSLLQSA   279
             * :*:****   :**.*********:*******.:*.******** .*:***********

mouse        WMEILILGIVYRSLPYDDKLAYAEDYIMDEEHSRLVGLLDLYRAILQLVRRYKKLKVEKE   339
human        WMEILILGIVYRSLPYDDKLVYAEDYIMDEEHSRLAGLLELYRAILQLVRRYKKLKVEKE   339
goat         WMEILILGIVYRSLPYDDKLVYAEDYIMDEEHSRLAGLLELYRAILQLVRRYKKLKVEKE   360
whale        WMEILILGIVYRSLPYDDKLVYAEDYIMDEEHSRLAGLLELYRAILQLVRRYKKLKVEKE   360
chicken      WMEILILGIVYRSLPYEDKLVYAEDYIMDEEHSRLTGLLELYLAILQLVRRYKKLKVEKE   360
fish         WMEILILSIVFRSLPYEDELVYAEDYIMDEEHSRLTGLLDLYVSILQLVRKYKKLKVEKE   339
             *******.**:*****:*:*.**************.***:** :******:*********

mouse        EFMILKALALANSDSMYIENLEAVQKLQDLLHEALQDYELSQRHEEPRRAGKLLLTLPLL   399
human        EFVTLKALALANSDSMYIEDLEAVQKLQDLLHEALQDYELSQRHEEPWRTGKLLLTLPLL   399
goat         EFVTLKALALANSDSMYIEDLEAVQKLQDLLHEALQDYELSQRHEEPRRTGKLLLTLPLL   420
whale        EFVTLKALALANSDSMYIEDLEAVQKLQDLLHEALQDYELSQRHEEPRRTGKLLLTLPLL   420
chicken      EFVTLKALALANSDSMHIEDMDAVQKLQDLLHEALQDYELSQRNEEPRRAGKLLLTLPLL   420
fish         EFVTLKAIALANSDSMHIEDMEAVQKLQDALHEALQDFECSQHQEDPRRAGKLLMTLPLL   399
             **: ***:********:**:::******* *******:* **::*:* *:****:*****

mouse        RQTAAKAVQHFYSVKLQGKVPMHKLFLEMLEAKV    433
human        RQTAAKAVQHFYSVKLQGKVPMHKLFLEMLEAKV    433
goat         RQTAAKAVQHFYSVKLQGKVPMHKLFLEMLEAKV    454
whale        RQTAAKAVQHFYSIKLQGKVPMHKLFLEMLEAKV    454
chicken      RQTAAKAVQHFYSIKLQGKVPMHKLFLEMLEAKV    454
fish         RQTATKAVQHFYSIKVQGKVPMHKLFLEMLEAKV    433
             ****:********:*:******************
```

##### 3.5.3. Disorder prediction

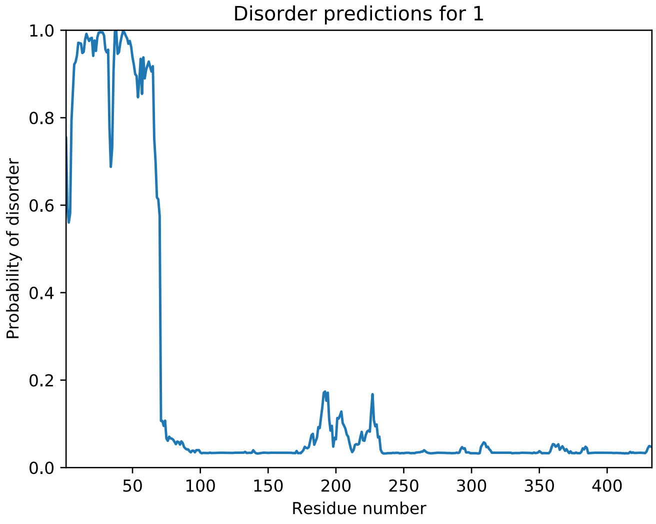


##### 3.5.4. Coding DNA sequences, produced protein constructs

**Esrrb-h_aa1-102_C12A-C72A-C91A**

Synthesized sequence:

```
Ccgcgggtgagaacctgtacttccagggcatgagcagcgaagatcgtcacctgggtagcagcgcgggcagctttattaaaaccgagccgagcag
cccgagcagcggtattgatgcgctgagccaccatagcccgagcggtagcagcgatgcgagcggtggcttcggtattgcgctgagcacccatgcg
aacggtctggatagcccgccgatgtttgcgggtgcgggcctgggtggcaacccggcgcgtaaaagctacgaagactgcaccagcggcatcatgg
aggatagcgcgattaaggcggaatatatgctgaacgcgattccgaaacgtctgtaaaagctt
```

Translates into:

```
AGENLYFQGMSSEDRHLGSSAGSFIKTEPSSPSSGIDALSHHSPSGSSDASGGFGIALSTHANGLDSPPMFAGAGLGGNPARKSYEDCTSGIME
DSAIKAEYMLNAIPKRL*
```

Expressed peptide:

GST-His-Tev-Esrrb(aa1-102)_C12A-C72A-C91A

```
MSPILGYWKIKGLVQPTRLLLEYLEEKYEEHLYERDEGDKWRNKKFELGLEFPNLPYYIDGDVKLTQSMAIIRYIADKHNMLGGCPKERAEISM
LEGAVLDIRYGVSRIAYSKDFETLKVDFLSKLPEMLKMFEDRLCHKTYLNGDHVTHPDFMLYDALDVVLYMDPMCLDAFPKLVCFKKRIEAIPQ
```

IDKYLKSSKYIAWPLQGWQATFGGGDHPPKSDGSTSGSGHHHHHHSAGENLYFQGMSSEDRHLGSSAGSFIKTEPSSPSSGIDALSHHSPSGSS
DASGGFGIALSTHANGLDSPPMFAGAGLGGNPARKSYEDCTSGIMEDSAIKAEYMLNAIPKRL

After Tev-cleavage *(leaving 1 Gly in N-ter):*

GMSSEDRHLGSSAGSFIKTEPSSPSSGIDALSHHSPSGSSDASGGFGIALSTHANGLDSPPMFAGAGLGGNPARKSYEDCTSGIMEDSAIKAEY
MLNAIPKRL

          10         20         30         40         50         60
G MSSEDRHLGS SAGSFIKTEP SSPSSGIDAL SHHSPSGSSD ASGGFGIALS THANGLDSPP

        70         80         90        100
MFAGAGLGGN PARKSYEDCT SGIMEDSAIK AEYMLNAIPK RL

#### Esrrb-h_aa1-102_C12A-C91A

Coding sequence (mutated from Esrrb-h aa-102 C12A-C72A-C91A):
Ccgcgggtgagaacctgtacttccagggcatgagcagcgaagatcgtcacctgggtagcagcgcgggcagctttattaaaaccgagccgagcag
cccgagcagcggtattgatgcgctgagccaccatagcccgagcggtagcagcgatgcgagcggtggcttcggtattgcgctgagcacccatgcg
aacggtctggatagcccgccgatgtttgcgggtgcgggcctgggtggcaacccgtgccgtaaaagctacgaagactgcaccagcggcatcatgg
aggatagcgcgattaaggcggaatatatgctgaacgcgattccgaaacgtctgtaaaagctt

Translates into:
AGENLYFQGMSSEDRHLGSSAGSFIKTEPSSPSSGIDALSHHSPSGSSDASGGFGIALSTHANGLDSPPMFAGAGLGGNPCRKSYEDCTSGIME
DSAIKAEYMLNAIPKRL*

After Tev-cleavage *(leaving 1 Gly in N-ter):*

GMSSEDRHLGSSAGSFIKTEPSSPSSGIDALSHHSPSGSSDASGGFGIALSTHANGLDSPPMFAGAGLGGNPCRKSYEDCTSGIMEDSAIKAEY
MLNAIPKRL

          10         20         30         40         50         60
G MSSEDRHLGS SAGSFIKTEP SSPSSGIDAL SHHSPSGSSD ASGGFGIALS THANGLDSPP

        70         80         90        100
MFAGAGLGGN PCRKSYEDCT SGIMEDSAIK AEYMLNAIPK RL

#### Esrrb-h_aa1-102_C12A-C72A-C91A_AviTag-His6

Synthesized sequence:
Ccgcgggtgagaacctgtacttccagggcatgagcagcgaagatcgtcacctgggtagcagcgcgggcagctttattaaaaccgagccgagcag
cccgagcagcggtattgatgcgctgagccaccatagcccgagcggtagcagcgatgcgagcggtggcttcggtattgcgctgagcacccatgcg
aacggtctggatagcccgccgatgtttgcgggtgcgggcctgggtggcaacccggcgcgtaaaagctacgaagactgcaccagcggcatcatgg
aggatagcgcgattaaggcggaatatatgctgaacgcgattccgaaacgtctgtaaaagctt

Translates into:
MSSEDRHLGSSAGSFIKTEPSSPSSGIDALSHHSPSGSSDASGGFGIALSTHANGLDSPPMFAGAGLGGNPARKSYEDCTSGIMEDSAIKAEYM
LNAIPKRLGLNDIFEAQKIEWHEGAGLE

Expressed peptide:
MSSEDRHLGSSAGSFIKTEPSSPSSGIDALSHHSPSGSSDASGGFGIALSTHANGLDSPPMFAGAGLGGNPARKSYEDCTSGIMEDSAIKAEYM
LNAIPKRLGLNDIFEAQKIEWHEGAGLEHHHHHH*

Expressed peptide:
MSSEDRHLGSSAGSFIKTEPSSPSSGIDALSHHSPSGSSDASGGFGIALSTHANGLDSPPMFAGAGLGGNPARKSYEDCTSGIMEDSAIKAEYM
LNAIPKRLGLNDIFEAQKIEWHEGAGLEHHHHHH

        10         20         30         40         50         60
MSSEDRHLGS SAGSFIKTEP SSPSSGIDAL SHHSPSGSSD ASGGFGIALS THANGLDSPP

        70         80         90        100
MFAGAGLGGN PARKSYEDCT SGIMEDSAIK AEYMLNAIPK RL GLNDIFEAQKIEWHEGAGLEHHHHHH

#### Esrrb-h_aa1-102_C12A-C72A_AviTag-His6

Coding sequence (mutated from Esrrb-h_aa-102_C12A-C72A-C91A_AviTag-His6):
Catatgagcagcgaagaccgtcacctgggtagcagcgcgggtagctttattaagaccgaaccgagcagcccgagcagcggcattgatgcgctga
gccatcatagcccgagcggtagcagcgatgcgagcggtggcttcggtattgcgctgagcacccatgcgaacggtctggatagcccgccgatgtt
tgcgggtgcgggcctgggtggcaacccggcgcgtaagagctacgaggactgcaccagcggcatcatggaggatagcgcgattaagtgcgaatat
atgctgaacgcgattccgaaacgcctgggcctgaacgacattttttgaagcgcagaagattgagtggcatgagggtgcgggcctcgag

Translates into:
MSSEDRHLGSSAGSFIKTEPSSPSSGIDALSHHSPSGSSDASGGFGIALSTHANGLDSPPMFAGAGLGGNPARKSYEDCTSGIMEDSAIKCEYM
LNAIPKRLGLNDIFEAQKIEWHEGAGLE

Expressed peptide:
MSSEDRHLGSSAGSFIKTEPSSPSSGIDALSHHSPSGSSDASGGFGIALSTHANGLDSPPMFAGAGLGGNPARKSYEDCTSGIMEDSAIKCEYM
LNAIPKRLGLNDIFEAQKIEWHEGAGLEHHHHHH

```
        10         20         30         40         50         60
MSSEDRHLGS SAGSFIKTEP SSPSSGIDAL SHHSPSGSSD ASGGFGIALS THANGLDSPP

        70         80         90        100
MFAGAGLGGN PARKSYEDCT SGIMEDSAIK CEYMLNAIPK RL GLNDIFEAQKIEWHEGAGLEHHHHHH
```

#### Esrrb-h_aa1-102_C12A-C72A-C91A-S22A_AviTag-His6

Coding sequence (mutated from Esrrb-h_aa-102_C12A-C72A-C91A_AviTag-His6):
Catatgagcagcgaagaccgtcacctgggtagcagcgcgggtagctttattaagaccgaaccgagcgcgccgagcagcggcattgatgcgctga
gccatcatagcccgagcggtagcagcgatgcgagcggtggcttcggtattgcgctgagcacccatgcgaacggtctggatagcccgccgatgtt
tgcgggtgcgggcctgggtggcaacccggcgcgtaagagctacgaggactgcaccagcggcatcatggaggatagcgcgattaaggcggaatat
atgctgaacgcgattccgaaacgcctgggcctgaacgacattttttgaagcgcagaagattgagtggcatgagggtgcgggcctcgag

Translates into:
MSSEDRHLGSSAGSFIKTEPSAPSSGIDALSHHSPSGSSDASGGFGIALSTHANGLDSPPMFAGAGLGGNPARKSYEDCTSGIMEDSAIKAEYM
LNAIPKRLGLNDIFEAQKIEWHEGAGLE

Expressed peptide:
MSSEDRHLGSSAGSFIKTEPSAPSSGIDALSHHSPSGSSDASGGFGIALSTHANGLDSPPMFAGAGLGGNPARKSYEDCTSGIMEDSAIKAEYM
LNAIPKRLGLNDIFEAQKIEWHEGAGLEHHHHHH*

```
        10         20         30         40         50         60
MSSEDRHLGS SAGSFIKTEP SAPSSGIDAL SHHSPSGSSD ASGGFGIALS THANGLDSPP

        70         80         90        100
MFAGAGLGGN PARKSYEDCT SGIMEDSAIK AEYMLNAIPK RL GLNDIFEAQKIEWHEGAGLEHHHHHH
```
